# Landscape drivers of genomic diversity and divergence in woodland Eucalyptus

**DOI:** 10.1101/590604

**Authors:** Kevin Murray, Jasmine Janes, Helen Bothwell, Ashley Jones, Rose Andrew, Justin Borevitz

**Affiliations:** Australian National University; University of New England

## Abstract

Spatial genetic patterns are influenced by numerous factors, and they can vary even among coexisting, closely related species due to differences in dispersal and selection. Eucalyptus (L’Héritier 1789; the “eucalypts”) are foundation tree species that provide essential habitat and modulate ecosystem services throughout Australia. Here we present a study of landscape genomic variation in two woodland eucalypt species, using whole genome sequencing of 388 individuals of *Eucalyptus albens* and *Eucalyptus sideroxylon*. We found exceptionally high genetic diversity (*π* ≈ 0.05) and low genome-wide, inter-specific differentiation (*F_ST_* = 0.15). We found no support for strong, discrete population structure, but found substantial support for isolation by geographic distance (IBD) in both species. Using generalised dissimilarity modelling, we identified additional isolation by environment (IBE). *Eucalyptus albens* showed moderate IBD, and environmental variables have a small but significant amount of additional predictive power (i.e., IBE). *Eucalyptus sideroxylon* showed much stronger IBD, and moderate IBE. These results highlight the vast adaptive potential of these species, and set the stage for testing evolutionary hypotheses of interspecific adaptive differentiation across environments.

## Introduction

In wild species, and especially plants, genetic variation is inherently spatial: individuals occur at specific locations, and allele frequencies differ across the landscape as a result of variation in demographic history, patterns of gene flow, and heterogeneous selection pressures. Landscape genomics is the study of the geographic distribution of alleles within a species and the underlying processes that shape gene flow. By interrogating spatial genetic patterns, we may examine the historical drivers of local genetic isolation and potential adaptation, and use this knowledge to better manage species under a changing environment (Hoffmann et al., 2015).

A multitude of processes may drive the spatial patterns of genetic diversity within and between species. Individuals may cluster into discrete genetic groups, with reduced gene flow between subpopulations relative to within. There are many potential causes of such discrete structure, for example geographic barriers to gene flow or flowering time divergence. Individuals may also exhibit patterns of continuous isolation by geographic distance (IBD; Wright, 1943) or isolation by environment (IBE; Wang and Bradburd, 2014). IBD is indicated by a positive correlation between increasing genetic dissimilarity and geographic distance, and is observed when individuals are more likely to reproduce with geographically-proximate individuals. IBE is indicated by a correlation between genetic dissimilarity and environmental dissimilarity, while controlling for IBD. IBE can have many causes, for example environmental effects on phenology altering flowering time, or impeded dispersal between habitats due to maladaptation to local conditions. Any of these three patterns of genetic isolation over the landscape (discrete structure, IBD, or IBE) may occur within a given species. Importantly, these patterns describe genome-wide phenomena, and while they may be influenced or initially generated by selection on adaptive alleles, their detection is not evidence of local adaptation. While factors affecting dispersal, such as landscape resistance (Spear et al., 2010; Wang and Bradburd, 2014; Zeller et al., 2012), may vary across the landscape, much can be learned by applying these global, homogeneous, dissimilarity-based methods for studying IBD and IBE, particularly when integrated with tests of discrete genetic structure.

The processes that influence spatial autocorrelation of allele frequencies require sophisticated statistical methods to disentangle. Continuous isolation by distance can lead to support for discrete population structure in analysis with genetic clustering methods like STRUCTURE and ADMIXTURE (Frantz et al., 2009). However, recent methodological developments now allow joint estimation of IBD and discrete structure (conStruct; Bradburd et al., 2017). Spatial autocorrelation of environmental variables makes disentangling their effects from IBD challenging, and older methods like partial Mantel tests are beset with several flaws (e.g., assumption of linearity, high Type I error rate; Guillot and Rousset, 2013). Generalised dissimilarity modelling (GDM; Ferrier et al., 2002, 2007) is a method which can accurately discriminate the geographic and environmental contributions to genetic differentiation, even where effects are non-linear. Equally important is the selection of variables appropriate to one’s study system: Williams et al. (2012) propose a comprehensive variable set and variable selection methodology specifically for ecological models of habitats. However sophisticated the methods used to detect isolation by environment, it is a pattern affecting the genomic background. Locally adaptive loci should stand out above this background and could be identified subsequently via a genome scan.

Genus *Eucalyptus* (L’Héritier; the “eucalypts”) is a speciose lineage of trees and large shrubs that includes the keystone species of many Australian habitats. Box-gum grassy woodlands are one such habitat, and while once common in southeastern Australia, their conversion to agricultural land has reduced their range significantly (NSW Scientific Committee, 2002). We sought to examine spatial genetic patterns in two foundation species of these grassy woodlands, *Eucalyptus albens* (Benth.; “white box”) and *Eucalyptus sideroxylon* (A. Cunn. ex Wools; “mugga ironbark”). The prevalence of discrete population structure, IBD and/or IBE has been studied in several eucalypt species (e.g. Andrew et al., 2005, 2007; Jones et al., 2007; Jordan et al., 2017; Rutherford et al., 2018; Steane et al., 2006, 2015, 2014; Supple et al., 2018). Although eucalypts have very limited seed dispersal, they generally preferentially outcross and are pollinated by generalist bird and insect pollinators, both of which contribute to their spatial genetic structure (Booth, 2017; Potts and Gore, 1995; Williams and Woinarski, 1997). Spatial genetic autocorrelation is strong within populations, but tends to be weak at larger scales; for example, isolation by distance between localities is only apparent between localities separated by more than 500 km in E. melliodora (Supple et al., 2018). While many studies have tested for and found discrete genetic structure (e.g. in *E. globulus*; Steane et al., 2006), strong discrete genetic structure uncorrelated with geography has been reported less commonly in widespread eucalypt species (e.g. in *E. salubris*; Steane et al., 2015). In any case, given the likely conflation of IBD and discrete population structure by traditional genetic clustering methods (Bradburd et al., 2017; Frantz et al., 2009), the relative extent of IBD and discrete structure remains an open question in many species. Correlation between genetic variation and environment has been observed in many forms, including IBE (e.g. Supple et al., 2018) and genotype-environment associations (e.g. Jordan et al., 2017; Dillon et al., 2014; Steane et al., 2017a, 2017b, 2014).

We aimed to determine the relative influence of the various factors contributing to landscape-scale spatial genetic patterns in *E. albens* and *E. sideroxylon*. The large estimated census sizes (González-Orozco et al., 2016) of both species led us to predict that these species would exhibit high genetic diversity. The reproductive ecology and extensive latitudinal geographic ranges of these species, and previous results for closely related species, led us to expect weak patterns of IBD and little discrete population structure orthogonal to IBD in both these species. Given gene-environment associations observed in closely-related species, we also predicted that isolation by environment would be observed, particularly associations between genetic distance and variables describing the availability of and demand for moisture and nutrients. To test these hypotheses, we generated whole-genome sequence data for 215 and 173 individuals of *E. albens* and *E. sideroxylon*, respectively. We quantified intraspecific genetic variation across the landscape, determined the extent of both continuous isolation by distance and isolation by environment, and assessed discrete population structure independent of IBD.

## Methods

### Study system

The genus *Eucalyptus* (L’Héritier 1789; the “eucalypts”) is described as a highly speciose lineage of trees and large shrubs within family *Myrtaceae*. Of the more than 800 described species (Nicolle, 2018; Pryor and Johnson, 1971) that have evolved over the last 70 My (Thornhill et al., 2015), nearly all are endemic to the Australian continent, with a small number of species occurring on tropical islands north of Australia. Here we focus on two woodland eucalypt species. *Eucalyptus albens* and *E. sideroxylon* are from different series (*Buxeales* and *Melliodorae*, respectively) within *Eucalyptus* section *Adnataria*. They are morphologically distinct, differing in bark type (box vs ironbark) and flower size and colour (*E. sideroxylon* larger, sometimes pink-red pigmented; Brooker and Kleinig, 2006; Boland et al., 2006; Costermans, 1983). Both generally occur inland of the Great Dividing Range, with *E. sideroxylon*’s range extending further inland, while *E. albens* extends further south and has disjunct populations in southeast Victoria and South Australia (see fig. 1). While both species have discontinuous distributions, partly as a result of post-European land clearing, *E. sideroxylon*’s distribution is believed to have been more discontinuous pre-colonisation (Costermans, 1983). Despite their largely sympatric distributions, there appears to be some niche specialisation between these species, with *E. albens* occupying more fertile soils, and *E. sideroxylon* preferring drier, well-drained, more gravelly soils (Boland et al., 2006; Costermans, 1983; Harden, 2000). Despite their classification into different series, there is evidence of ongoing gene flow between these species, with reports of hybrid zones (Pryor, 1953), as is common in *Eucalyptus* generally, and especially in section Adnataria (Griffin et al., 1988).

**Figure 1:**
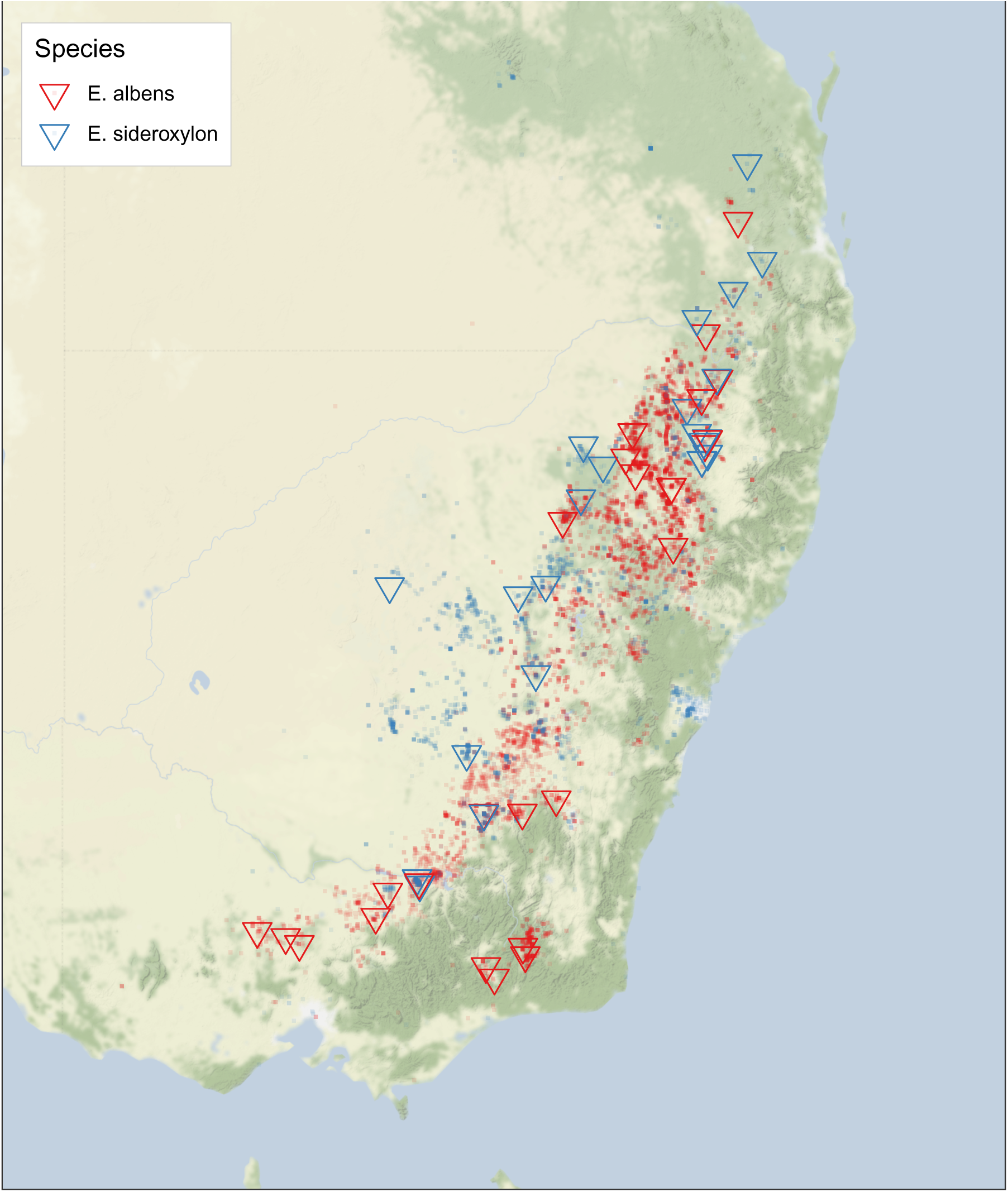
Focal species occurrence records and sampling localities. Geolocated occurrence records (± km accuracy) for *E. albens* and *E. sideroxylon* obtained from the Atlas of Living Australia are overlain on a map of southeastern Australia. Sampling localities used in this study are indicated by large triangles.

### Data acquisition

Samples used in this study were collected from naturally occurring trees of the target species throughout southeastern Australia. Leaf tissue and fruit were collected from between 3 and 15 trees from each location, across 39 distinct locations (fig. 1). Sample identifiers, GPS locations, and additional metadata are presented online (https://doi.org/10.6084/m9.figshare.7583291.v1). Sampling was performed between 2015 and 2017, primarily by Dr Jasmine Janes. Leaves were dried on silica gel, and 20-30 3 mm leaf hole punches were taken for DNA extraction (Harris Uni-Core WB100039). Hole punches were added to 1.1 mL mini-tubes (Axygen Scientific) with a 3mm ball bearing, frozen under liquid nitrogen, and ground for 2 min using a TissueLyser (Qiagen). DNA extraction was performed using a 96-well column based kit, Invisorb DNA Plant HTS 96 Kit/ C 96 well purifications (Stratec Molecular 7037300400). The protocol was performed following the manufacturer’s instructions, except for the lysis incubation, which was extended from 1 hour to 2 hours.

Multiplexed, short-read, whole-genome shotgun DNA sequencing libraries were generated using a cost-optimised, transposase-based protocol (Jones et al., 2018). Briefly, fluorometric DNA quantification was performed using a Quant-iT^TM^ high sensitivity dsDNA assay kit (Molecular Probes^TM^ Q33120). DNA was diluted to 2 ng*/µ*L, quantified again and then diluted to 0.8 ng*/µ*L, normalising concentrations across all samples. Then, 3 *µ*L of each sample (2.24 ng) was transferred to a new plate with a small quantity of a Nextera^TM^ tagment DNA enzyme (Illumina catalogue #15027865) to add adapters (tagmentation). This reaction was optimised to be 1/25th of manufacturer’s protocol, to save reagents and increase throughput. Custom index primers were used to amplify the libraries during 13 cycles of PCR (primer sequences provided in Jones et al., 2018). Libraries were purified and size-selected with a combination of bead- and electrophoresis-based methods, selecting fragments with insert sizes between 200 and 500 bp. These purified libraries were sequenced on a variety of Illumina platforms, with most libraries sequenced on multiple runs across both NextSeq 500 and NovoSeq 2000 instruments at the Biomolecular Resource Facility, ANU, and the Ramaciotti Center, UNSW. Multiple runs were pooled by sample to obtain sufficient coverage. Library preparation was performed primarily by Dr Ashley Jones and Dr Norman Warthmann.

### Alignment and polymorphism detection

Sequencing yielded between 3 Gbp and 10 Gbp per sample, pooled across all sequencing runs (see fig. 2). Raw sequence data was quality filtered using AdapterRemoval (Schubert et al., 2016), removing adaptor sequences, trimming low-quality (<Q25) sub-sequences, and merging overlapping read pairs. We used BWA MEM version 0.7.15 (Li, 2013; Li and Durbin, 2009) to align short reads using default alignment parameters to the *Eucalyptus grandis* reference genome, with an assembled *E. grandis* chloroplast added to the nuclear genome assembly (HM347959; Paiva et al., 2011). Across all samples, 90% percent of reads were aligned to the *E. grandis* reference, with an average alignment mismatch rate of 4.8%. Both read mapping and alignment mismatch rates suggest a reference bias between species (with *E. sideroxylon* appearing less distant).

**Figure 2:**
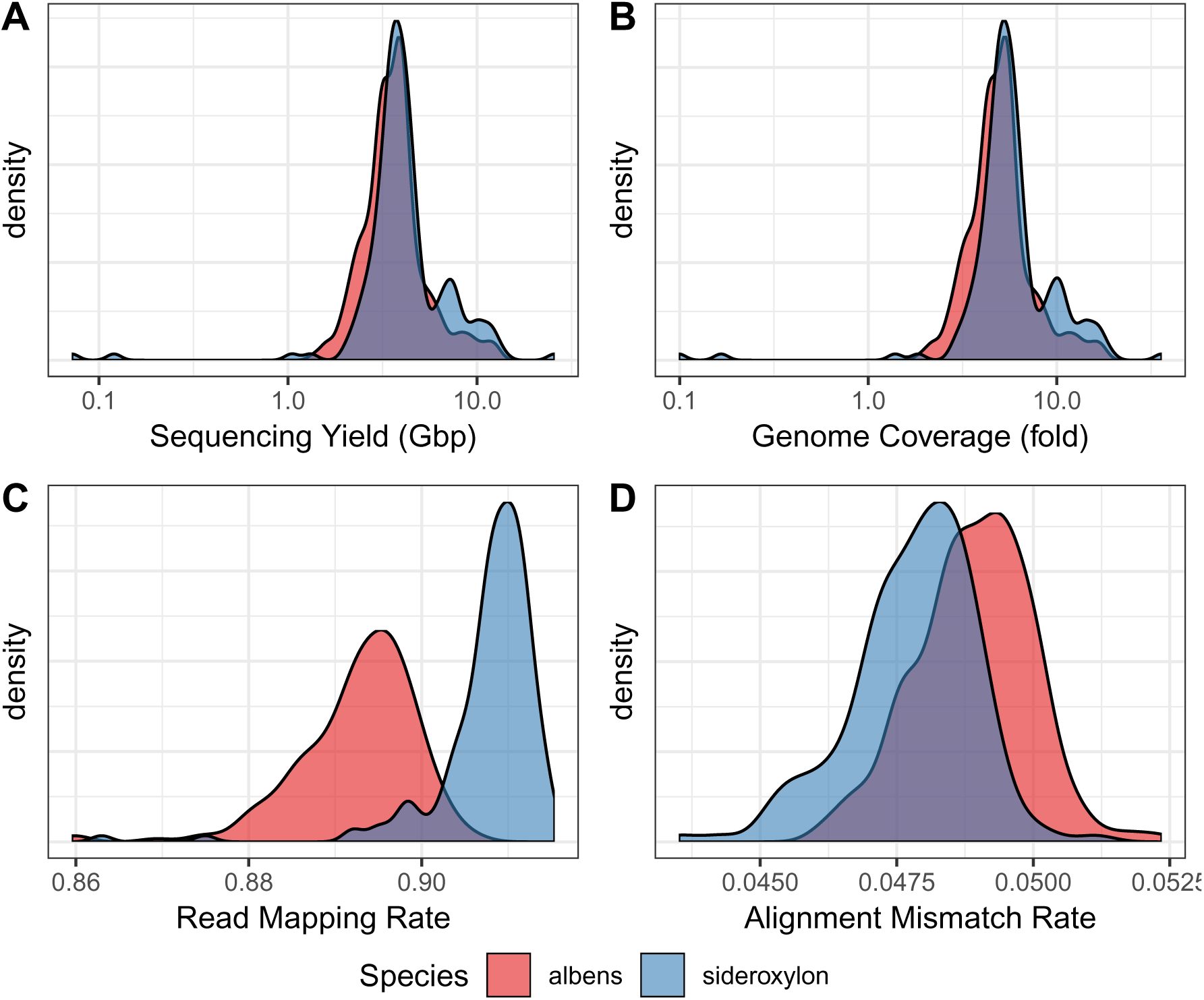
Whole genome sequencing yield and alignment statistics. A) Post-QC raw sequencing yield in bases, showing most samples yielded between 3 Gbp and 10 Gbp. B) *E. grandis* genome coverage (total sum of aligned bases). C) Read alignment rate, D), proportion of aligned bases which do not match the *E. grandis* reference genome. Overall, we have consistent, moderate coverage (median 5.2-fold), although both read mapping and alignment mismatch rates suggest a reference bias between species (with *E. sideroxylon* appearing less distant).

We detected short genomic variants using an efficient pipeline implementing the variant calling models contained in FreeBayes (Garrison and Marth, 2012) and `bcftools mpileup` (Li, 2011). As these tools are not internally parallelised, and the volume of data generated in this project was very large, I developed a genomic region-parallelised system pipeline around these software. Briefly, this pipeline performs variant calling on each 100 kbp region of the *E. grandis* reference genome in parallel across hundreds of CPUs at once, before merging the candidate variants discovered in each region into a genome-wide variant set. This variant set was then normalised with `bcftools norm` (Li, 2011), block substitutions were decomposed to single nucleotide polymorphisms (SNPs) using `vt decompose_blocksub` (Tan et al., 2015), and filtered with `bcftools filter`. We discarded variants with quality less than 10, fewer than five reads in total across all alleles in all samples, and fewer than three reads supporting the alternate allele across all samples. In total, we discovered 132 million putative variants, of which 55 million were common (*>*10% minor allele frequency) SNPs within at least one species.

While many analyses require knowledge of exact genotypes for each sample, some methods (e.g. ANGSD; Korneliussen et al., 2014) are able to represent uncertainty in individual genotypes through subsequent analyses. Given our low sequencing coverage, individual genotypes may have higher error than we desire, particularly in detecting heterozygosity. To address these concerns, we used ANGSD (Korneliussen et al., 2014) to detect putative variants, and to calculate genotype likelihoods at each variable site. ANGSD considered loci only if there were *>*10 reads at a SNP (summeds across all samples), considered reads only if they had a mapping quality *>*30, considered bases within reads only if they had a base quality score *>*10, and removed variants with a minor allele frequency <2%, with fewer than three reads supporting the alternate allele, or if the p-value of the likelihood-ratio test of non-zero minor allele frequency (i.e. test of polymorphism) was *>*10*^−^*^3^. Indel and block-substitution variation is not considered by ANGSD. We used a region-parallel approach similar to that used in variant calling to accelerate this computation. In total, ANGSD detected 55 million polymorphisms across our samples.

From ANGSD likelihoods, we calculated several population genetic statistics. A two-dimensional site-frequency spectrum (SFS) between all *E. albens* and *E. sideroxylon* was calculated with realSFS (Nielsen et al., 2012), then estimated genome-wide *F_ST_* between *E. albens* and *E. sideroxylon* using this two-dimensional SFS as a prior (see Supplementary fig. 15). Using ngsDist (Fumagalli et al., 2014), we calculated inter-sample genetic distances for all non-outlier samples. We estimated inter-sample covariance using PCAngsd (Meisner and Albrechtsen, 2018). We calculated Euclidean distances from PCAngsd covariances using the Gower transformation (*D_ij_* = *C_ii_* + *C_jj_* − 2*C_ij_*; Gower, 1985).

All steps in the above pipeline have been implemented as a generic, modular Snakemake workflow. In particular, the region-parallelisation of variant calling is handled specifically in Snakemake, allowing abstraction of the execution environment. Project and cluster specific configuration of this pipeline is separate to pipeline code, allowing easy adaptation to other systems and datasets. In fact, this pipeline has subsequently been used in at least three additional projects (wheat, tomato, and potato population genomics). This pipeline and associated scripts are open source, and available online at https://github.com/kdmurray91/euc-dp14-workspace.

### Population genetic analysis

We performed kmer-based exploratory genetic analysis, to confirm sample identities and guide subsequent analyses. Genetic distances were estimated using kWIP, a kmer-based estimator of genetic distance (Murray et al., 2017). We first counted 21-mers in unaligned, quality trimmed sequencing reads, after pooling all reads for each sample into one file. We estimated inter-sample genetic distances using the weighted inner product metric implemented in kWIP. Distances were estimated on each data subset (all 10 Adnataria species, both *E. albens* and *E. sideroxylon*, and *E. albens* and *E. sideroxylon* separately) to allow subset-specific weighting. We visualised these exploratory analyses using both hierarchical clustering (`hclust`) and classical multidimensional scaling (`cmdscale`) in R 3.4 (R Core Team, 2018). In addition to kmer-based estimates of genetic distance, we visualised the sample covariance (or genomic relationship matrix) as estimated by PCAngsd in a similar fashion, and compared these results visually.

To examine within-locality diversity, a variety of population diversity metrics were employed. We calculated Nei’s sample-size corrected gene diversity (or expected heterozygosity, 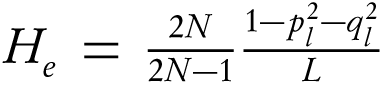; Nei and Roychoudhury, 1974), using per-locality allele frequencies calculated from expected genotypes by PCAngsd. Additionally, we calculated gene diversity for all pairs of sampling locations, by considering all individuals from both pairs as a single site (equivalent to gene diversity in a sample-size weighted mean of allele frequencies). We displayed these measures of intra- and inter-location genetic diversity by plotting location estimates on a map of south-eastern Australia using ggmap and Stamen map layers (Kahle and Wickham, 2013).

Traditional model-based genetic clustering methods like STRUCTURE (Pritchard et al., 2000) and ADMIXTURE (Alexander et al., 2009) were designed to detect discrete population structure, therefore they may perform poorly for continuously distributed natural populations in which isolation by distance is the primary driver of genetic structure (Frantz et al., 2009). ConStruct addresses this limitation by jointly modeling the effects of both continuous isolation by distance and discrete population structure on inter-sample relationships (Bradburd et al., 2017). As we expected continuously distributed landscape features to contribute to inter-sample genetic distances, we used conStruct to simultaneously test for discrete and continuous population structure. We used per-locality allele frequencies calculated from PCAngsd expected genotypes. We tested two distinct models separately for *E. albens* and *E. sideroxylon*, using the cross-validation approach implemented in conStruct: a model similar to that used by STRUCTURE, and one allowing for isolation by distance within genetic clusters (‘layers’). Layer contributions were calculated for all cross-validation runs. To test for recent admixture between *E. sideroxylon* and *E. albens*, we used conStruct directly on the estimated genotypes, again performing cross-validation and calculating layer contributions.

We estimated the distribution of genome-wide linkage disequilibrium by calculating inter-SNP correlations and modeling correlation decay as a function of chromosomal position. Using the BoringLD R package (https://github.com/kdmurray91/boringld), we first calculated pairwise *r*^2^ among SNPs in 30 kbp genomic windows with an overlap of 10 kbp between adjacent windows from FreeBayes-called variants. Then, we fitted analytical models of the decay of *r*^2^ as a function of inter-SNP base pair distance to determine the recombination parameter (*ρ*), using formulae derived by Hill and Weir (1988). The base pair distance to half-maximal *r*^2^ was also calculated for each window. Window estimates of both *ρ* and half-maximal *r*^2^ were summarised across all genome windows.

### Landscape genomic analyses

We used Generalised Dissimilarity Modelling (GDM) to test for isolation by distance without assuming a linear relationship between geographic and genetic distance. Using genetic distances derived from PCAngsd covariance, we modeled genetic distance as a function of geographic distance within each species. We calculated geographic distances between samples with earth.dist from the `fossil` R package (Vavrek, 2011). Models were constructed using individual-level genetic and geographic distances, using three I-spline knots. Only distance pairs with a geographic distance greater than 10 kilometers (i.e. inter-location pairs) were considered. For each model, we examined the robustness of spline fits using jackknifing with 100 replicates. For each jackknife replicate, we removed all samples from a random 10% of sampling locations and fitted the GDM models as before. To perform cross-validation of each model, we partitioned data into training and test sets comprising 90% and 10% of sampling locations, respectively. We then computed cross-validation accuracy as the correlation between actual genetic distances for all distances for the 10% test data partition, and the corresponding distances predicted using a GDM model trained on samples from the 90% training data partition.

To assess isolation by environment, we first selected potentially relevant environmental variables based on a general methodology described by Williams et al. (2012). Variable values were extracted using the Atlas of Living Australia’s (ALA) Spatial Portal (“Atlas of Living Australia,” 2018). To determine which variables to include in models of IBE, we first performed forward selection within each category: Water, Energy, and Soil (see Supplementary tbl. 2). We excluded terrain and geoscientific variables, as these processes vary over finer spatial scales than our aggregated sampling resolution. In each forward selection run, we started with a GDM model of genetic distance as a function of geographic distance, and proceeded by adding the variable that, when included, increased the proportion of deviance explained by the model by the largest amount. We terminated this process when no variable could explain at least 1% of additional deviance. We then combined forward-selected variables across all categories into a candidate GDM model. To assess how representative our sampling was of each species’ range, we compared distributions of each environmental variable from our sampling locations to distributions for ALA observation records for each species.

To refine candidate GDM models, and assess the importance and significance of constituent variables, we performed backward selection using the `gdm.VarImp` function in the GDM package (Ferrier et al., 2007; Manion et al., 2018), with 100 permutation replicates for each step. For both species, the inflection point in decreased model deviance explained resulted in five variables retained for the final model (Supplementary fig. 12). We then assessed the consistency of spline fits using the jack-knifing approach described above. These new functions for variable selection and cross-validation are available as an R package (https://github.com/kdmurray91/gdmhelpers).

## Results

### Population genetic variation

After filtering unsupported or singleton variants, we discovered over 100 million candidate variants (varying slightly between software tools; Supplementary tbl. 1). This equates to about 1/6th of all positions in the *E. grandis* reference genome. Of these candidate variants, around 40% were not segregating (<10% minor allele frequency) in either *E. albens* and *E. sideroxylon*. Of the remaining approximately 60 million variants, over half were segregating in both species, with 22% private to *E. albens* and 23% private to *E. sideroxylon* (Supplementary tbl. 1). ANGSD estimated inter-species genome-wide *F_ST_* between *E. albens* and *E. sideroxylon* to be 0.15; global intraspecific *F_ST_* was 0.018 in *E. albens* and 0.017 in *E. sideroxylon*.

Using kmer-based estimators of genetic distance, we estimated genome-wide differentiation within the Adnataria. A principal component analysis (PCA) on kWIP distance estimates showed four clusters corresponding to taxonomic series. In some cases, species formed discrete subgroups within series, though in many cases species clusters overlapped somewhat (fig. 3); within-series divergence between species varies. Hierarchical clustering of kWIP distances showed similar patterns. The two focal species of this study formed clearly distinct clusters, as expected (Supplementary fig. 13).

**Figure 3:**
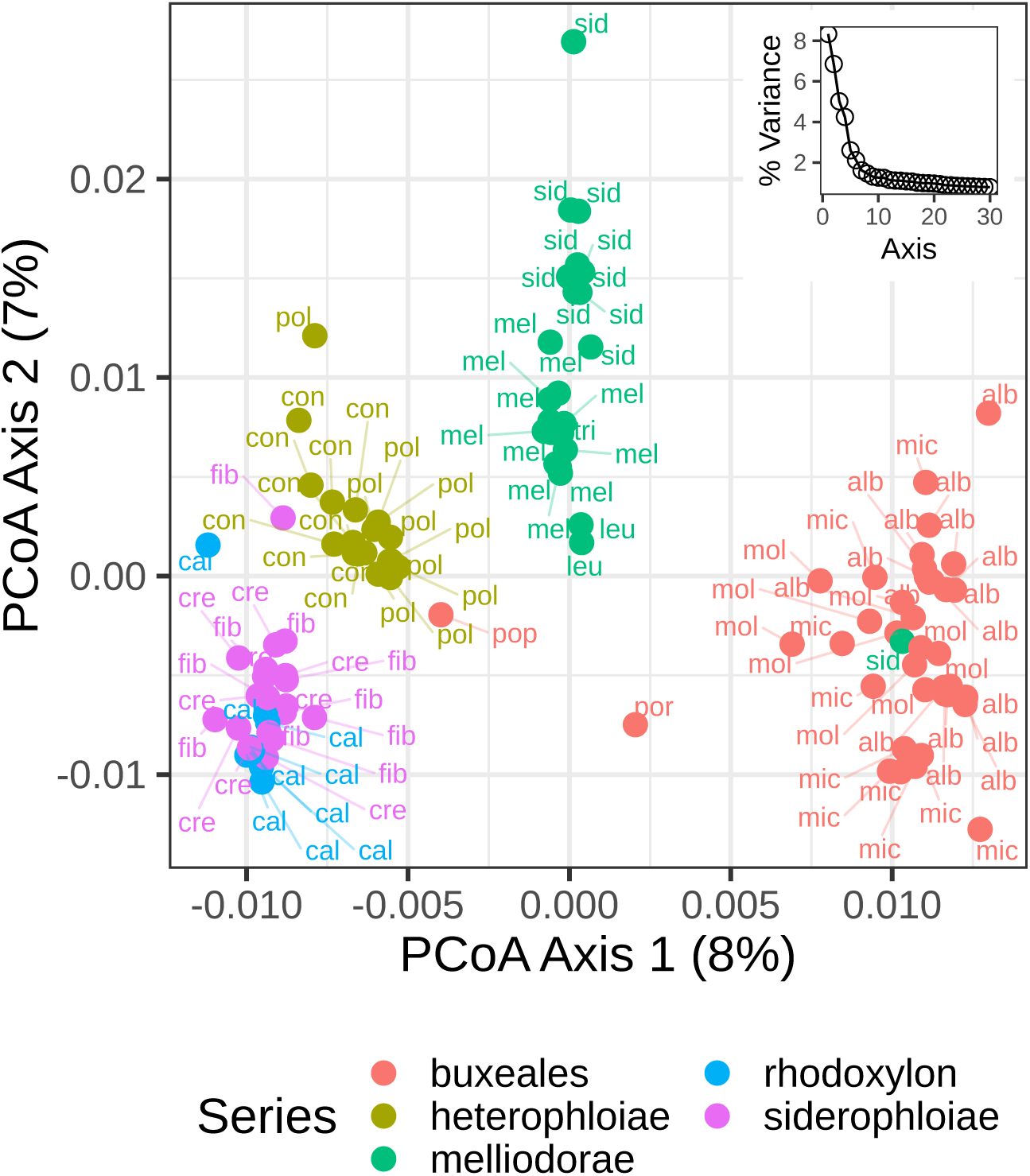
Intra- and inter-series genomic divergence. Principal coordinates analysis was performed on distances calculated using kWIP directly on short reads without alignment to a reference. Broadly, all samples from across five series in Adnataria form four clusters, corresponding to series-level divergences. Two series form a single cluster, Rhodoxlyon, and Siderophloiae; recent taxonomies reclassify *E. caleyi* within Siderophloiae (Nicolle, 2018). Within-series divergence between species varies. Individuals’ species are denoted using the first three letters of species names: alb - *E. albens*, cal - *E. caleyi*, con - *E. conica*, cre - *E. crebra*, fib - *E. fibrosa*, leu - *E. leucoxylon*, mel - *E. melliodora*, mic - *E. microcarpa*, mol - *E. mollucana*, pol - *E. polyanthemos*, pop - *E. populnea*, sid - *E. sideroxylon*, tri - *E. tricarpa*. Samples appearing far from their species/series clusters likely represent either misidentification or sample mislabels, and were excluded from subsequent analyses.

*Eucalyptus albens* and *E. sideroxylon* had high genetic diversity. Expected heterozygosity within sampling locations ranged between 0.2 and 0.3 for both species, with *E. sideroxylon* having slightly lower mean location-level diversity, particularly in northern localities. Both species exhibited high species-wide genetic diversity (*E. sideroxylon H_e_* = 0.25, *π* = 0.053; *E. albens H_e_* = 0.26, *π* = 0.056). Background linkage disequilibrium (LD) decayed rapidly in both species (Supplementary fig. 11). The median base-pair distance to half-maximal *r*^2^ in *E. albens* was 92 bp (IQR 47-219 bp), while LD extended slightly further in *E. sideroxylon* (median 113 bp; IQR 55-264 bp).

### Spatial genetic diversity and structure

In general, genetic diversity was spread evenly over the range of our sampling in both species (fig. 4). Both *π* and *H_e_* are almost equal across all locations sampled in *E. albens*, while genetic diversity in *E. sideroxylon* declined very slightly in locations toward the north of our sampling. Similarly, expected heterozygosity and *π* among samples at pairs of locations were uncorrelated with pairwise geographic distance (Supplementary fig. 14).

**Figure 4:**
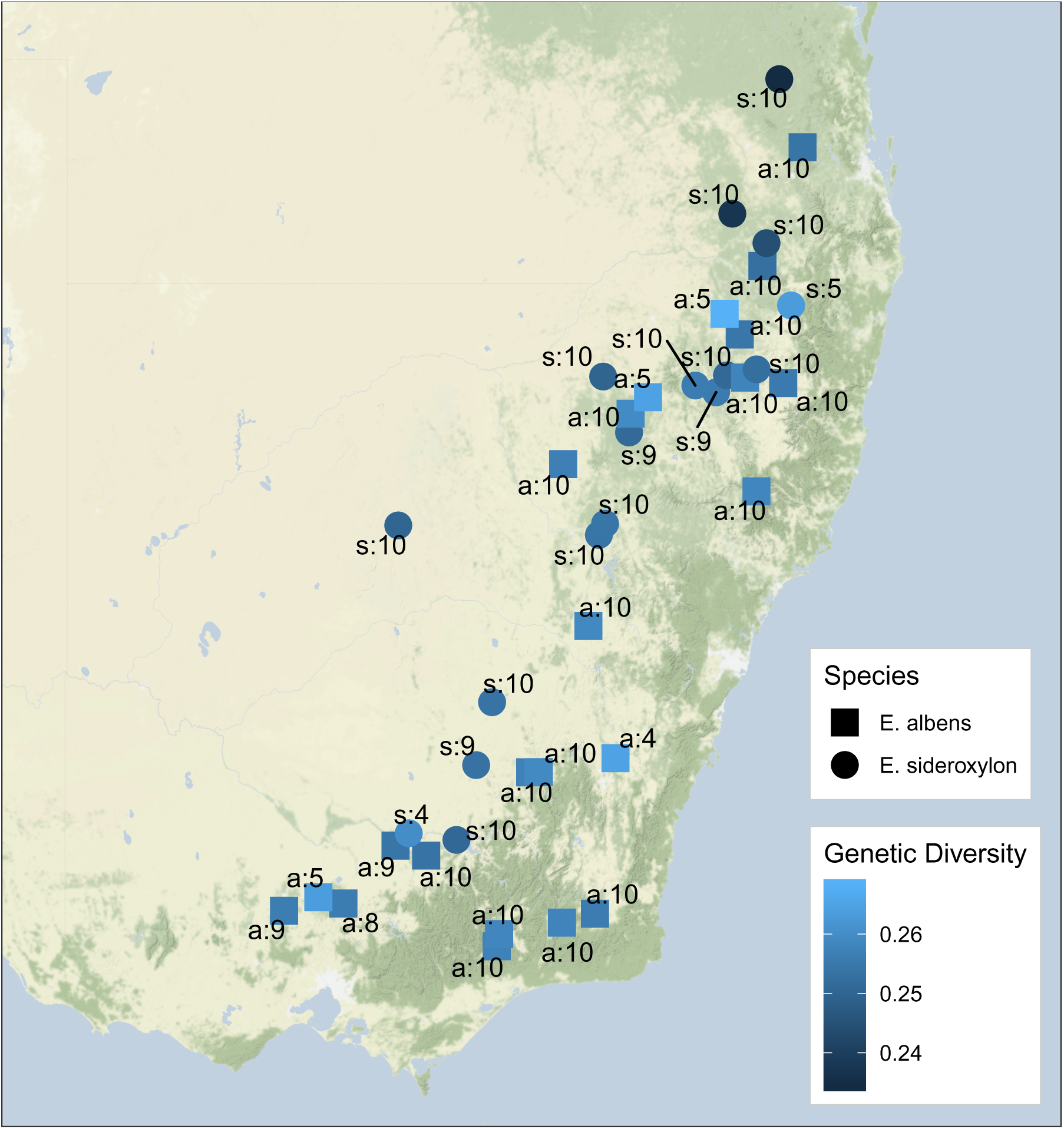
Geographic surface of genetic diversity superimposed on a map of southeastern Australia. Annotations describe species (s: or a: for *E. sideroxylon* and *E. albens* respectively) and the number of individuals per locality.

### No discrete but continuous population structure

Neither *E. albens* or *E. sideroxylon* exhibited strong signs of discrete population structure in a PCA of intra-sample genetic covariance as estimated by PCAngsd (fig. 5). Leading principal component axes explained little of the overall genomic variance between samples (0.8% and 0.6% in *E. albens*, 3.6% and 1.0% in *E. sideroxylon*). In each species, the leading principal component axis was correlated with latitude, suggesting isolation by geographic distance (*E. albens r*^2^ = 0.92, *p* < 0.0001; *E. sideroxylon r*^2^ = 0.87, *p* < 0.0001).

**Figure 5:**
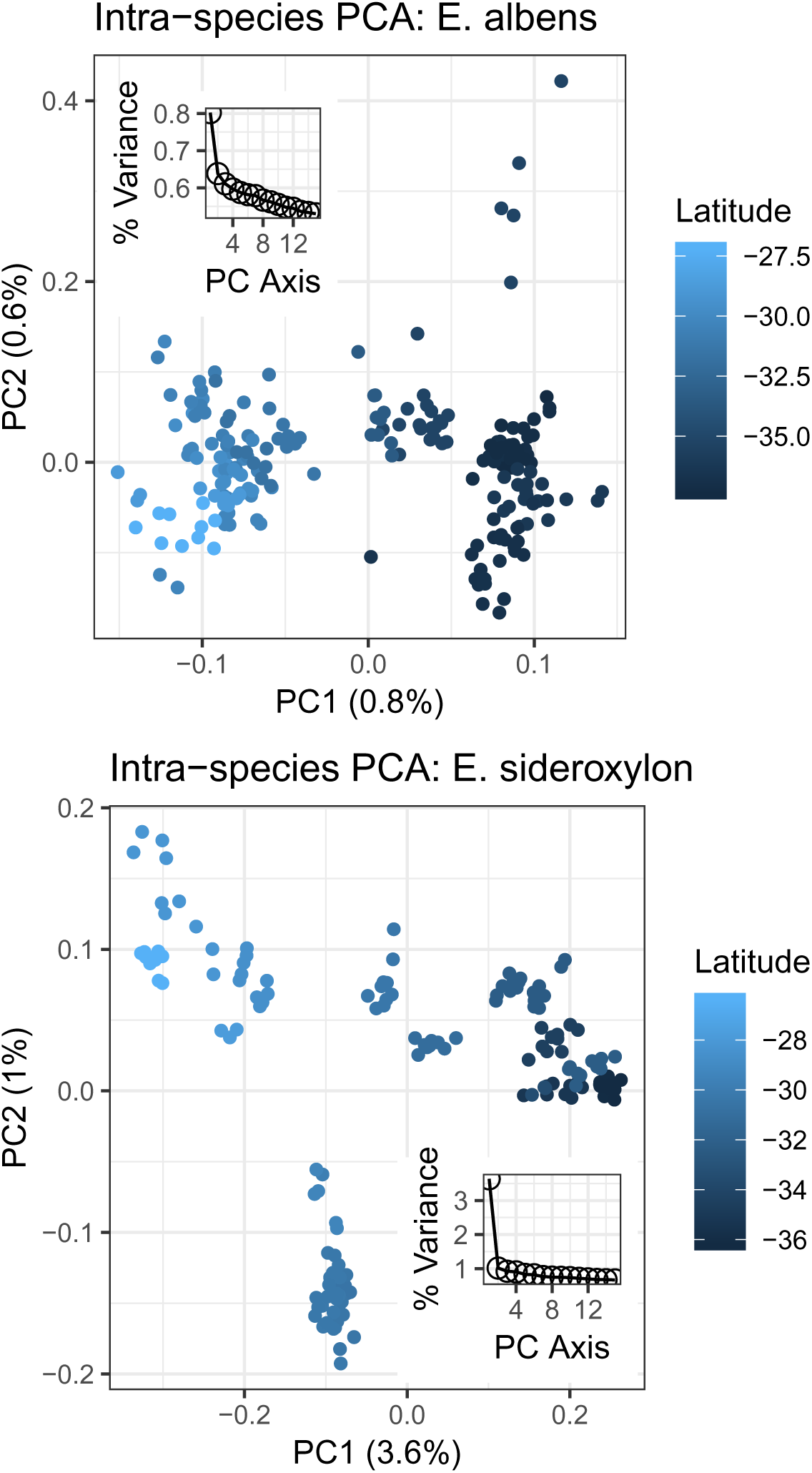
Principal component analysis (PCA) of *E. albens* and *E. sideroxylon* individual genotypes. Axes describe eigendecomposition of PCAngsd estimates of sample covariance. Individuals are coloured by latitude, the primary axis of variation in species’ distributions. Insets show the distribution of leading eigenvalues. Note the absence of strong discrete clusters, the strong trend in PC1 across latitude, and the low proportion of genetic variance explained by each leading axis.

Joint estimation of continuous isolation by distance and discrete population structure indicated both species likely form single, continuous populations, with clinal structure influenced by strong IBD. When accounting for IBD in conStruct, cross-validation of conStruct models suggested either one or two populations in both species (fig. 6). In models with two population layers, the second layer contributed very little additional predictive accuracy. The second layer in such models had no strong signal of IBD. This second layer could describe a small contribution of inter-species introgression to extant genetic diversity, or could represent “homogeneous minimum layer membership”, an artifact produced by conStruct when there are significant levels of missing data (Bradburd et al., 2017). ConStruct models that did not allow continuous isolation by distance required at least two populations to achieve similar predictive accuracy (fig. 6).

**Figure 6:**
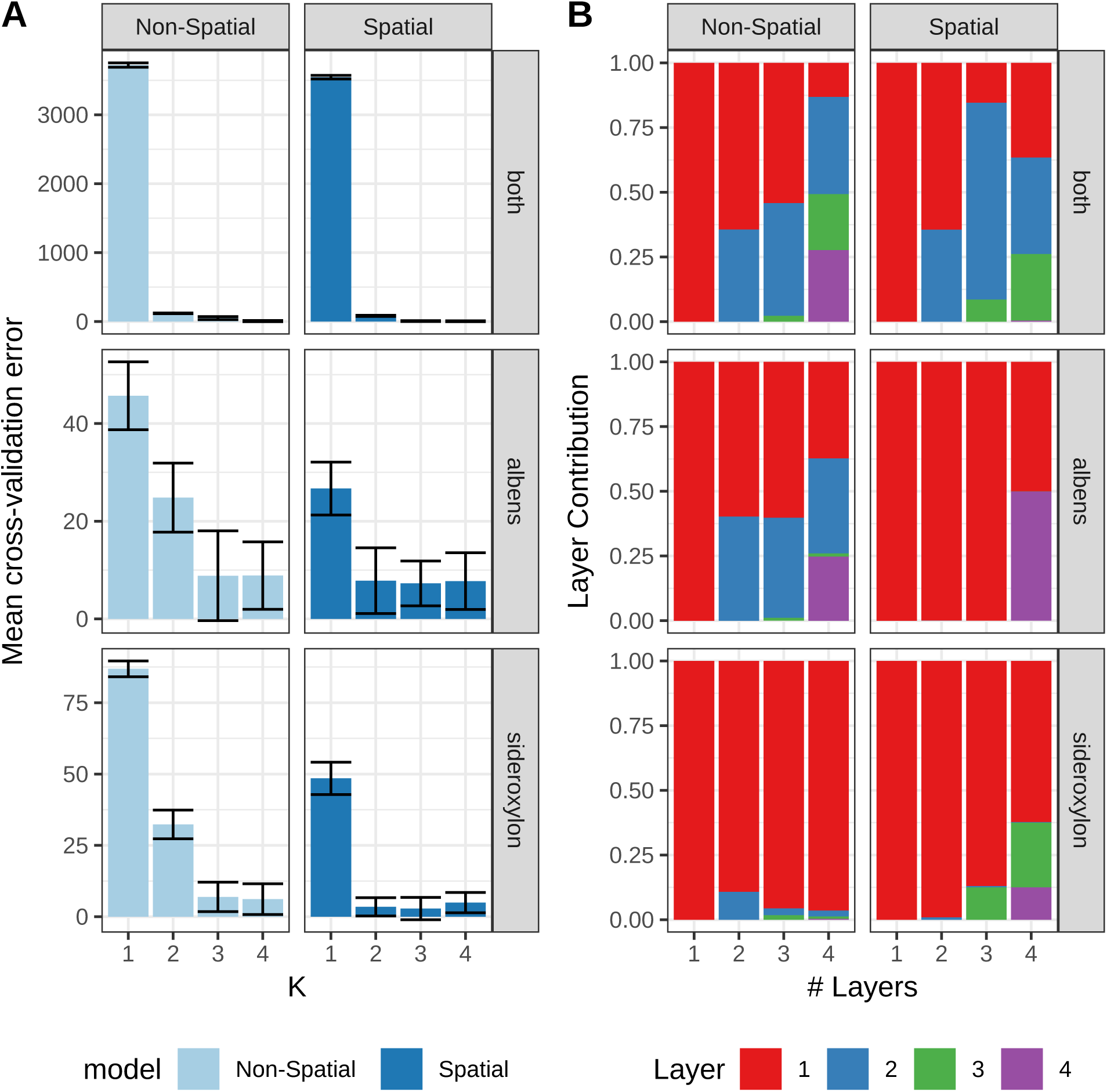
Cross-validation of conStruct models of continuous and discrete population structure. A) Model cross-validation error, means ± SD. B) Layer contribution to model explanatory power within each model with “# Layers”. Non-spatial: construct models that do not account for IBD, Spatial: construct models that allow for IBD within each population layer. Plots rows are for datasets with all localities across both *E. albens* and *E. sideroxylon* (“both”) or within each species.

### Interspecific gene flow

We detected signals suggesting ongoing inter-species gene flow. Six samples were intermediate between *E. albens* and *E. sideroxylon*, being both intermediate in PCA (Supplementary fig. 13), and having interspecies admixture proportions between 30% and 70% (Supplementary fig. 16). Two of these samples were identified as putative hybrids in the field. Mantel tests of inter-species distance pairs showed weak but statistically significant correlation between genetic distance and geographic distance, indicating that co-located *E. albens* and *E. sideroxylon* had lower genetic distance than geographically distant samples. This pattern could be due to inter-series gene flow, and is not predicted by incomplete lineage sorting, but could also be caused by certain demographic histories (e.g., expansion from shared ancestral refugia). Individual admixture proportions estimated by conStruct models supported the status of these six samples as recent hybrids (Supplementary fig. 16). Additionally, conStruct models suggested a variable, small proportion (between 0% and 10%; Supplementary fig. 16) of admixture from *E. albens* to *E. sideroxylon* (or vice versa). Additionally, more than half of all variants that were common in either species were common in both species (Supplementary tbl. 1). These results concur with ABBA-BABA-based formal tests of admixture conducted in an as-yet-unpublished sister study (J. Janes, pers. comm.).

### Isolation by distance and environment

Isolation by distance was moderately strong and largely linear in both species. Using generalised dissimilarity modelling (GDM) to model genetic distance as a function of geographic distance, we found *E. albens* to have moderately strong, almost linear IBD, with models explaining approximately 26% of overall deviance (*P* < 0.001; fig. 7). Meanwhile, *E. sideroxylon* exhibited very strong IBD, with models explaining 78% of overall deviance (*P* < 0.001; fig. 7). The relationships described by the best fit splines were robust to the removal of 10% of the sampling locations (i.e. jackknifing; fig. 7).

**Figure 7:**
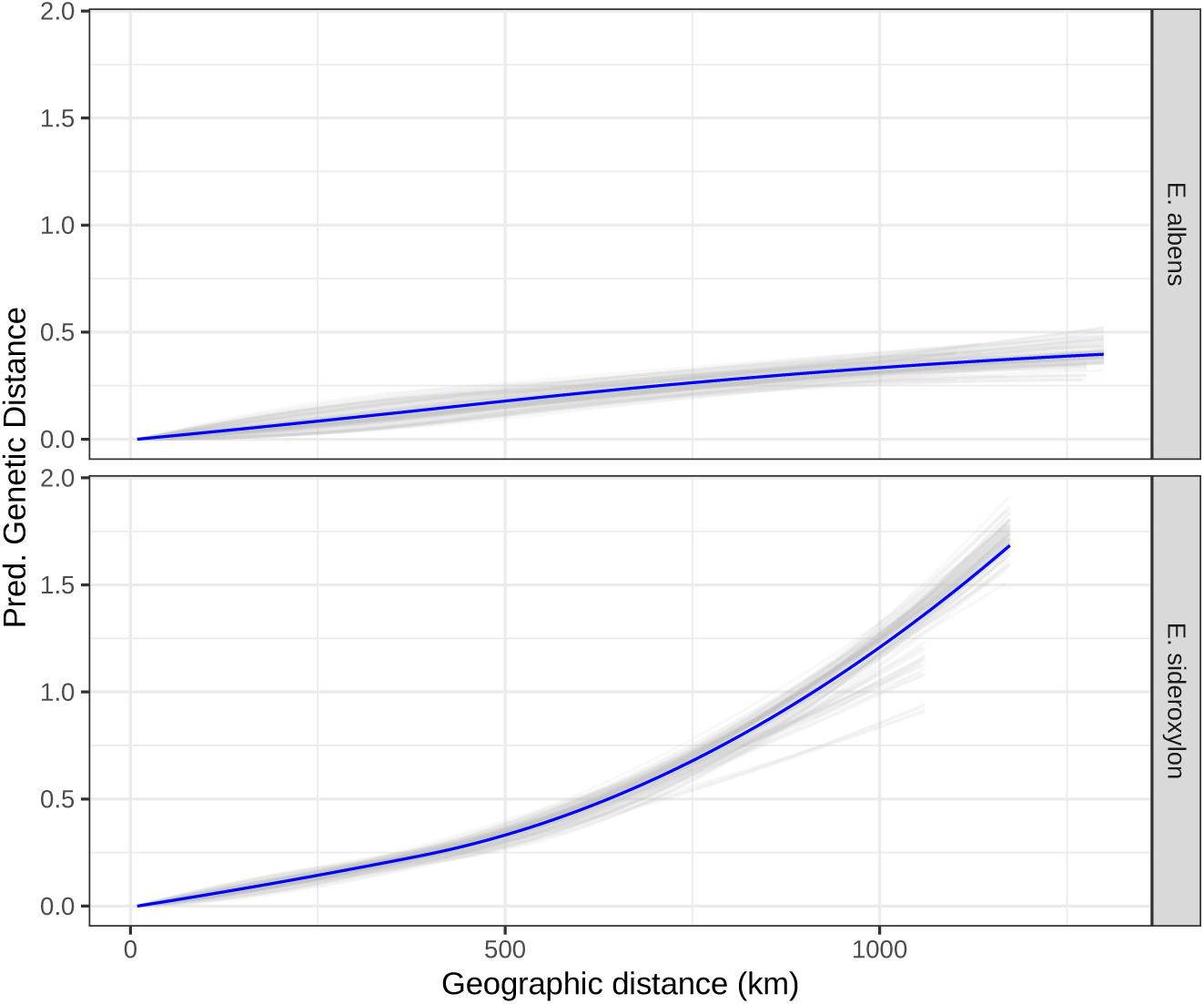
Geographic GDM show strong isolation by distance. GDM model splines (blue) and jackknife replicate splines (grey) that best describe the association between geographic distance and genetic distance in each species. Geography-only GDM models explain 26% of model deviance in *E. albens*, and 78% in *E. sideroxylon*. IBD appears to have an approximately linear trend in *E. albens*, while the strength of IBD increases for *E. sideroxylon* localities separated by more than 500 km.

In the GDM analysis with environmental predictors, *E. albens* showed moderate isolation by environment, particularly driven by precipitation and substrate related environmental variables. Forward selection identified 11 candidate environmental covariates, each able to explain at least 1% additional deviance. Backward selection on these 11 variables identified substrate hydrological conductivity, substrate phosphorus concentration, spring/autumn precipitation seasonality, precipitation of the wettest quarter, and total wind run as contributing the highest predictive power (Supplementary tbl. 3). Overall, this model explained 31% of total deviance (*P* < 0.001), 7% higher than a model containing only geographic distance. Cross-validation showed this model to have reasonable predictive accuracy; the correlation between predicted and true genetic distances was *r*^2^ = 0.33, roughly equal to the percentage of deviance explained (Supplementary fig. 10). For most variables, splines of best fit were robust to removal of 10% of sampling locations, although some variables had high uncertainty (e.g. precipitation of the wettest month), and other variables showed bimodal distributions of spline fits (e.g. autumn/spring precipitation seasonality; fig. 8).

**Figure 8:**
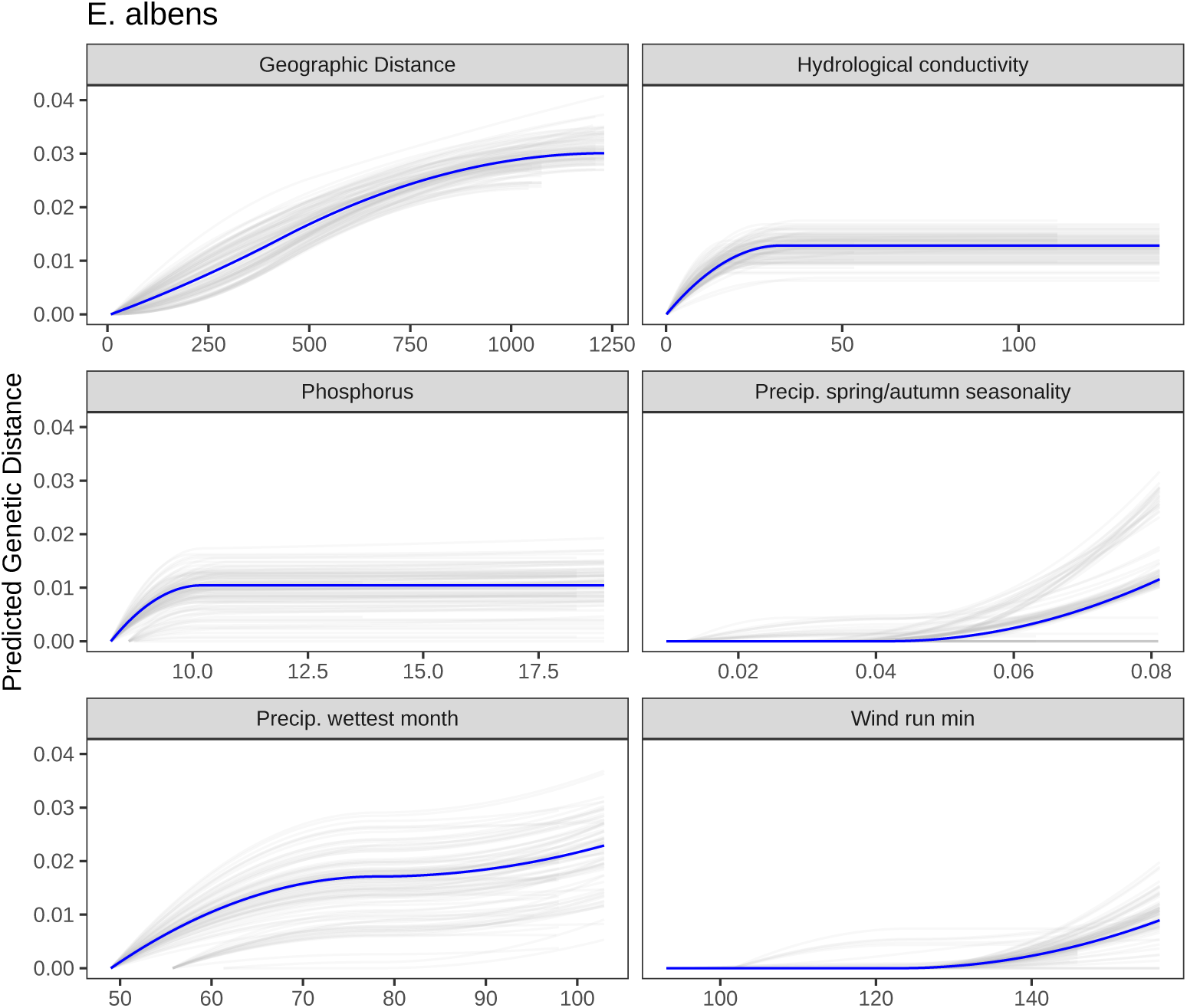
GDM spline fits for *E. albens*. To test the robustness of GDM predictive splines, models were re-run with 10% of sampling locations removed in each dataset. Each panel showed the range of spline fits among the 100 jackknife replicates (grey) and the full data (blue).

Similarly, *E. sideroxylon* showed somewhat stronger isolation by environment than *E. albens*, primarily driven by environmental variables describing the timing, availability, and demand for moisture. Forward selection identified 12 candidate covariates, and backward selection identified maximum cloud-adjusted solar radiation, maximum month-on-month differences in temperature and precipitation, maximal vapour pressure deficit, and substrate water holding capacity as the five variables with highest predictive power (Supplementary tbl. 3). Again, the overall model was highly significant (*P* < 0.001), explained 90% of total deviance (12% higher than a model containing only geographic distance), and had very high mean cross-validation predictive accuracy (*r*^2^ = 0.90; Supplementary fig. 10). Splines of best fit were robust to removal of 10% of sampling locations for all predictors, with low uncertainty in spline fits across jack-knifing replicates fig. 9.

**Figure 9:**
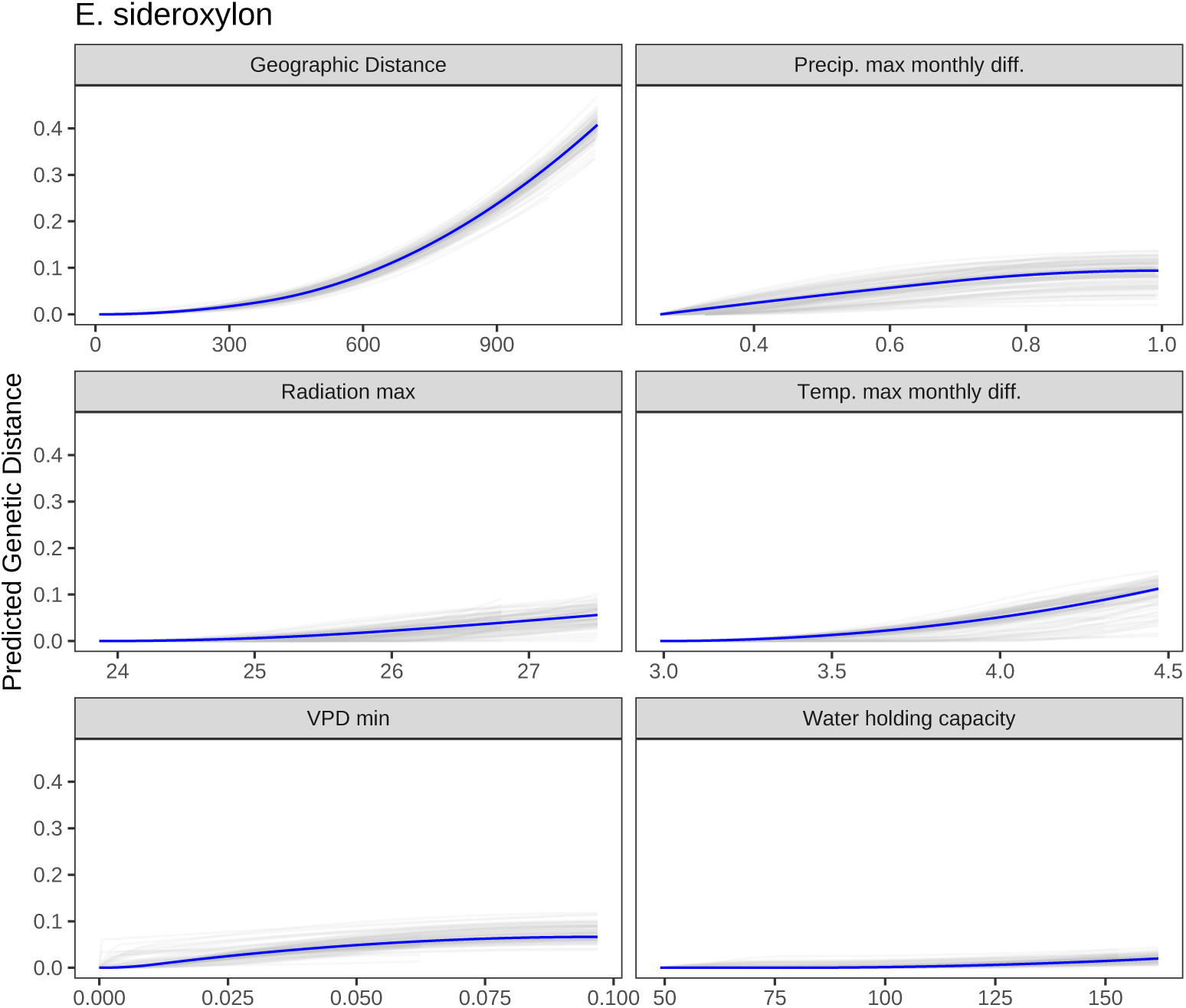
GDM spline fits for *E. sideroxylon*. To test the robustness of GDM predictive splines, models were re-run with 10% of sampling locations removed in each dataset. Each panel showed the range of spline fits among the 100 jackknife replicates (grey) and the full data (blue).

## Discussion

### Genetic diversity

Eucalypts form large, continuous populations with high genetic diversity and low population divergence. We confirm this result with one of the first whole-genome population resequencing studies in wild eucalypts (Kainer et al., 2018; Silva-Junior and Grattapaglia, 2015). We estimated intra-species *F_ST_* to be 0.017-0.018, lower than estimates from previous studies in a variety of eucalypt species (*E. melliodora*: *F_ST_* = 0.04, Supple et al., 2018; *E. globulus*: *F_ST_* = 0.08, Jones et al., 2002). These previous estimates are of similar magnitude to widespread tree species in other biomes, for example Oaks, Poplar, and Pine (*Quercus robur*: *F_ST_* = 0.07, Vakkari et al., 2006; *Q. engelmannii*: *F_ST_* = 0.04, Ortego et al., 2012; *Populus temuloidies*: *F_ST_* = 0.03, Wyman et al., 2003; *Pinus taeda*: *F_ST_* = 0.04, Eckert et al., 2010; *P. contorta*: *F_ST_* = 0.02, Yang et al., 1996). This very weak genetic structure likely results from a combination of very large, stable effective population sizes, widespread ranges, and high outcrossing rates (Williams and Woinarski, 1997).

While high compared to many tree species, genetic diversity both across all individuals and within localities is slightly lower in *E. sideroxylon* than *E. albens*. Previous work indicated especially high allozyme diversity in *E. albens* (Prober and Brown, 1994). Estimates of effective population size within *E. albens* and *E. sideroxylon* follow a similar pattern (J. Janes *et al*., in prep.). Linkage disequilibrium reported here is less extensive than in some previous reports (Silva-Junior and Grattapaglia, 2015), and is more similar to older estimates of LD decay from wild individuals of *E. grandis* (Grattapaglia and Kirst, 2008) and *E. globulus* (Thavamanikumar et al., 2011). A crucial caveat to these results is that we predominantly sampled from mature trees which likely predate the extensive land clearing and habitat fragmentation that accompanied European colonisation of Australia. The applicability of these results and conclusions to future generations of these species is uncertain. Individuals from later generations show reduced but still high genetic and/or phenotypic diversity in recent studies of related *Eucalyptus* species (Broadhurst, 2013; Jordan et al., 2016; Supple et al., 2018), although these studies examined planted individuals, either in provenance trials or revegetation efforts (Costa e Silva et al., 2011). Further research on the differences in genetic diversity between remnant stands and younger cohorts is warranted.

### Continuous genetic divergence

We observed continuous differentiation across the landscape within both species, driven both by geography and environment. This matches findings in most previous studies of genomic variation in eucalypts (Jordan et al., 2017; Steane et al., 2015, 2014; Supple et al., 2018). However, unlike previous studies, we found no support for strong discrete genetic structure. As seen in simulated and empirical studies of continuously distributed species (Bradburd et al., 2017; Frantz et al., 2009), we found statistical support for discrete population structure only when IBD was not incorporated into models of population structure. This conflation of IBD and discrete structure cements the conclusion that accurate determination of population structure in widespread species should use methods that can jointly estimate isolation by distance and discrete population structure.

We found very strong isolation by distance, particularly in *E. sideroxylon*. This is much stronger than in previous studies on related species at similar spatial scales. For example, weak isolation by distance occurs among populations in *E. melliodora*, with little correlation of genetic and geographic distance between pairs separated by less than 500 km (Supple et al., 2018; but see Andrew et al., 2005), and relatively weak IBD has been found in *E. microcarpa* (Jordan et al., 2017). Weak IBD may have technical and/or biological causes. Noisy reduced-representation sequencing methods that have large error in estimating sample genotypes (e.g. in *E. melliodora*; Supple et al., 2018), and therefore genetic distances, may have led to underestimation of the correlation between genetic and geographic distances. The difference in resolution in the present study may be partly due to our use of PCAngsd to calculate genetic distances, as it is designed to reduce the stochastic effects of low-coverage sequencing on inter-individual distances. The use of PCAngsd here is likely to have a similar effect to the PCA-based genetic distances recommended by Shirk et al. (2017).

Strong IBD is a result of by patterns of migration imposed by the reproductive ecology of eucalypts (Williams and Woinarski, 1997). Seed dispersal is limited in eucalypts, with pollen exchange accounting for the vast majority of migration among localities (Booth, 2017; Potts and Gore, 1995; Williams and Woinarski, 1997). Pollination is facilitated by generalist insect, bird, and mammal pollinators in nearly all species (Potts and Gore, 1995; Williams and Woinarski, 1997). Most exchanges of pollen occur within a limited local range; however, migration events occur over much longer ranges with lower frequency (Williams and Woinarski, 1997). As a result, genes are readily exchanged far beyond immediate neighbours. We found the strength of IBD to be strikingly different between *E. sideroxylon* and *E. albens*. This finding suggests that, while pollen-mediated gene flow is strong enough to limit discrete population structure in both species, gene flow at larger spatial scales is more restricted in *E. sideroxylon* than in *E. albens*. This goes against the expectation that the larger, more coloured flowers of *E. sideroxylon* attract more frequent bird pollination, leading to higher pollen motility. These observations are also supported by lower local genetic diversity within *E. sideroxylon*, particularly in northern localities.

### Isolation by Environment

We observed isolation by environment in both species, primarily driven by variables describing the availability of water and nutrients to plants, with little influence of temperature. Permutation-based variable testing showed only a small orthogonal contribution of environment to observed genetic distances, after accounting for geographic distance. Strong spatial autocorrelation of environment variables prevents fully disentangling geographic and environmental contributions to gene flow across the landscape. Exclusion of relevant environmental variables could cause underestimation of overall IBE, although the variable selection procedure employed here tested the contribution of a broad range of environmental variables concerning soil, geology, precipitation, temperature, wind, solar radiation, and aridity. In most cases, inference of the environmental drivers of genomic differentiation appear robust to subsampling of localities. GDM models of isolation by distance and environment had high cross-validation accuracy, and all were significant under locality-wise permutation testing. While specific environmental variables selected as most important were not shared, the strength of IBE was similar in both species. Furthermore, the variables most predictive of genetic distance in both species described the availability and demand for moisture or soil fertility (nutrient or water availability). Despite local niche separation (Boland et al., 2006; Brooker and Kleinig, 2006; Costermans, 1983; Harden, 2000), the ranges of *E. albens* and *E. sideroxylon* overlap significantly (fig. 1), and therefore likely experience selection along similar macro-scale clines (e.g. temperature, aridity).

Correlation of genetic and environmental variation is well established in Eucalyptus. Differences in climate and soil nitrogen can predict genetic differentiation in *E. melliodora* (Supple et al., 2018). Allele frequencies at certain SNPs were significantly correlated with aridity, temperature, and rainfall in *E. tricarpa* (Steane et al., 2014), *E. loxophleba* (Steane et al., 2017a), and *E. microcarpa* (Jordan et al., 2017). Our use of environmental variables designed to interrogate the ecology of Australian plants (Williams et al., 2012) precludes direct comparison of IBE among studies at the level of specific variables. However, our results follow a similar general pattern to these previous studies of gene-environment association in eucalypts.

### Interspecific Divergence and Gene Flow

About half of all common variants discovered in this study are common in both species, and we observed low genome-wide divergence between *E. albens* and *E. sideroxylon* (*F_ST_* = 0.15). Recent evidence suggests the genetic divergence is not strong at most genomic loci in many species, both in eucalypts (Rutherford et al., 2018) and more broadly (Andrew and Rieseberg, 2013; Wu, 2001). Additionally, low interspecific differentiation is expected theoretically given extremely large effective population sizes, long generation times, and relatively recent radiation (González-Orozco et al., 2016).

Interspecific gene flow between eucalypts has been observed many times, though probably occurs at a low rate in nature (Griffin et al., 1988). We made several observations suggestive of ongoing gene flow between *E. albens* and *E. sideroxylon*. We identified several putative hybrid individuals in the field, via PCA, and conStruct indicated a low but consistent proportion of inter-series admixture. Hybridisation between *E. albens* and *E. sideroxylon* has been demonstrated previously (Pryor, 1953), and more broadly, a systematic review by Griffin et al. (1988) showed species within Eucalyptus section Adnataria were found to hybridise at the highest rate of any section. The proportion of hybrids we observe here is of the same approximate magnitude as that observed in several other eucalypts in the subgenus *Symphomyrtus* (1-3%; Williams and Woinarski, 1997). Hybridisation between *E. albens* and *E. sideroxylon* occurs in spite of ecological differentiation, for example, in the form of limited local co-occurrence, different tolerance of poor soils and aridity (Boland et al., 2006; Costermans, 1983; Harden, 2000), and relatively little overlap in flowering period (*E. albens*: January-June, *E. sideroxylon* May-November; Costermans, 1983; Brooker and Kleinig, 2006).

### Conservation implications

To avoid extirpation, organisms must either adapt or migrate as environments change (Aitken et al., 2008). Our findings of high genetic diversity imply a large pool of variation accessible to natural selection. However, the long generation time of these trees makes it unlikely that natural selection on local standing variation alone can outpace anthropogenic changes in climate and land use; therefore, migration of better-adapted alleles is required (Booth, 2017; Booth et al., 2015). While we show pollen must have been exchanged over relatively large distances at a rate historically sufficient to prevent strong differentiation between localities, natural rates of migration seed-based migration are unlikely to be prevent range contractions (Booth, 2017; Prober et al., 2015). Human assistance may be required to shift the ranges of these and many other woodland species (Butt et al., 2013; González-Orozco et al., 2016; Supple et al., 2018).

Management interventions can take numerous forms. There is a temptation to use models of isolation by environment to guide selection of seed sources for assisted migration. However, we urge the utmost caution when doing so: these models of IBE are based on genome-wide patterns among predominantly near-neutral genetic variation, and use predicted, interpolated environmental data. Such models could detect the historical influence of environment on genetic diversity, but there is no promise that these influences reflect what may happen in the future. In particular, we strongly discourage the use of these results (or the results of any similar study) to narrow the range of seed sources used to revegetate any given locality. Studies of inbreeding and outbreeding depression in interspecific crosses of eucalypts find strong effects of selfing and local inbreeding (Hardner and Potts, 1995), but little difference in fitness proxies beyond hundreds of meters (Hardner et al., 1998). Such results reinforce the need for a restoration strategy that focuses on adaptive potential as much as pre-adapted germplasm. Our advice matches that proposed in numerous recent syntheses of revegetation strategy (Broadhurst et al., 2008; Kardos and Shafer, 2018; Prober et al., 2015; Weeks et al., 2011), in particular “climate-adjusted provenancing” (Prober et al., 2015). As an additional consideration, climate change is not the only anthropogenic risk to these species: the habitat these species inhabit has been cleared extensively since European colonisation of Australia, with only a few percent of the habitat remaining (NSW Scientific Committee, 2002). Perhaps the most effective management action would be the prevention of further deforestation and habitat fragmentation, both for these species and generally.

### Future directions

All patterns reported here concern genome-wide average effects; significant variation between loci in patterns described here likely exists. Investigating how variation in ancestry, population structure, interspecific differentiation, and associations with environment differ across the genome requires whole-genome datasets, and the dataset and analysis pipeline we present here enables these analyses. In particular, our finding of low linkage disequilibrium implies that many reduced-representation sequencing methods would provide data for just a fraction of all independent loci, and therefore miss important segregating variation (Ahrens et al., 2018; Lowry et al., 2017).

Genotype-environment association (GEA) studies could detect individual alleles which vary in frequency across some environmental cline, accounting for geography and genome-wide patterns (as has been observed with reduced representation sequencing in related species, e.g. Steane et al., 2014, 2017a, 2017b). Loci that have undergone selective sweeps could also be detected, shedding further light on recent evolution (Nielsen et al., 2005). Similarly, investigation of inter-species divergence at specific loci could highlight which loci are maintaining species boundaries in the face of gene flow (Strasburg et al., 2012). Finally, genome-wide average ancestry may differ significantly from local ancestry at nearly all loci across the genome, and could be examined in these species (e.g. using Local PCA; Li and Ralph, 2018).

## Conclusions

In summary, we found high intraspecific genetic diversity, low genome-wide divergence between *E. albens* and *E. sideroxylon*, and evidence of ongoing gene flow between these species. We found no evidence of strong, discrete population structure, and uncovered strong continuous isolation by distance in both species. We also found that isolation by geographic distance accounts for most, but not all, of this continuous genetic structure, with environmental variables describing the availability and demand for moisture, temperature, and substrate contributing to the pattern of IBE. Taken together, these results describe *E. albens* and *E. sideroxylon* as widespread species with high genetic diversity and strong isolation by distance. A small proportion of genetic variation is associated with climate; however, high levels of genetic diversity exist regionally, and even within localities. This high genetic diversity implies these species have high adaptive potential, especially if enhanced by assisted migration. The crucial test of these species’ survival will not be the level of understanding we gain about the intricacies of isolation by landscape, but rather the extent to which we utilise these and other species in large-scale rehabilitation of degraded ecosystems.

## Acknowledgements

We thank Norman Warthmann, Tim Collins, Jamieson Gorrell, Jeremy Bruhl, and Allison Huesler for technical assistance. This work was supported financially by the Australian Research Council (CE140100008; DP150103591), and an Australian Government Research Training Program scholarship. The research was undertaken with the assistance of resources from the National Computational Infrastructure (NCI), which is supported by the Australian Government.

## Supplementary information

**Table 1:**
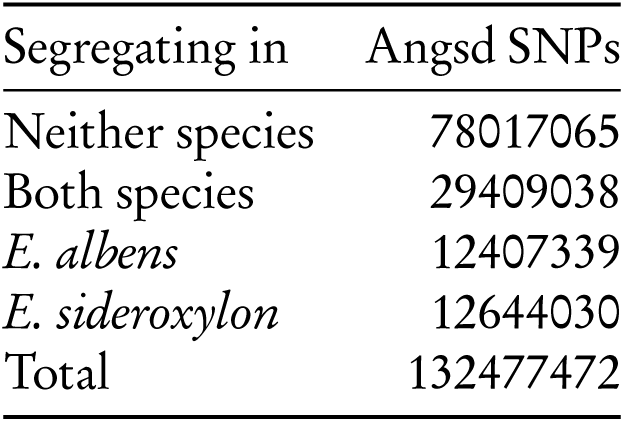
SNP genotyping statistics.

| Segregating in | Angsd SNPs |
| --- | --- |
| Neither species | 78017065 |
| Both species | 29409038 |
| *E. albens* | 12407339 |
| *E. sideroxylon* | 12644030 |
| Total | 132477472 |

**Table 2:**
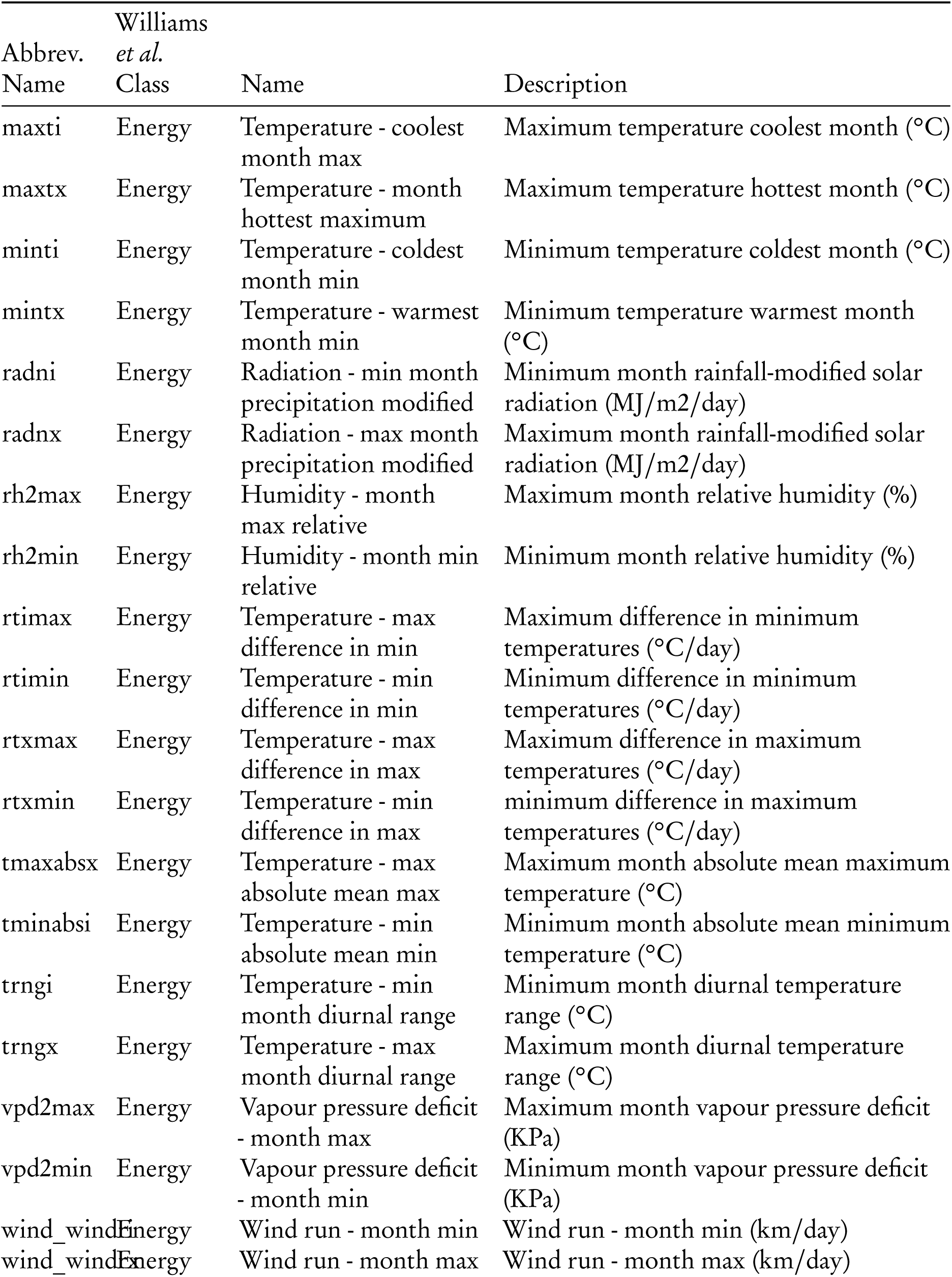
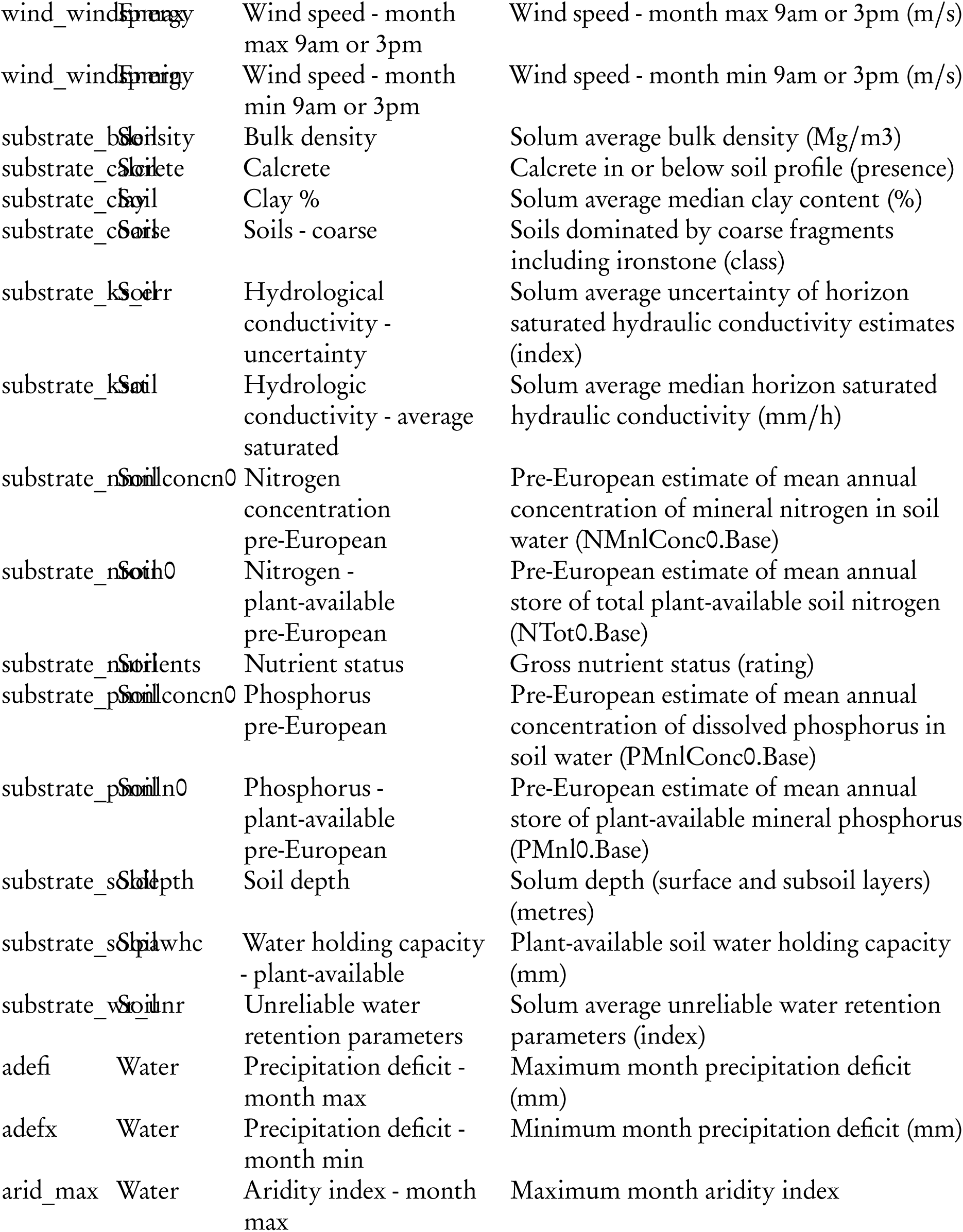
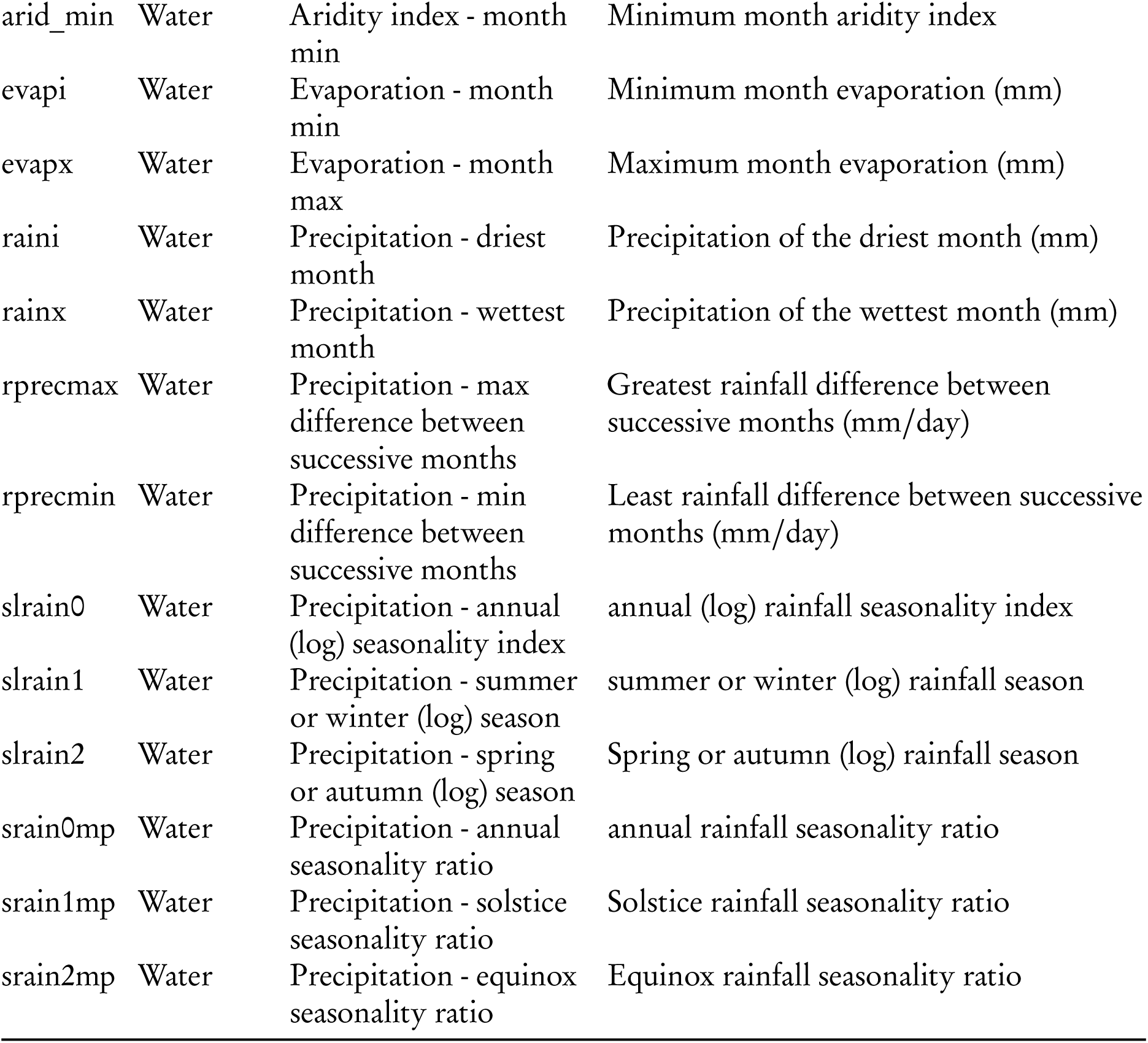
Environmental variables considered in forward selection of IBE models.

**Figure 10:**
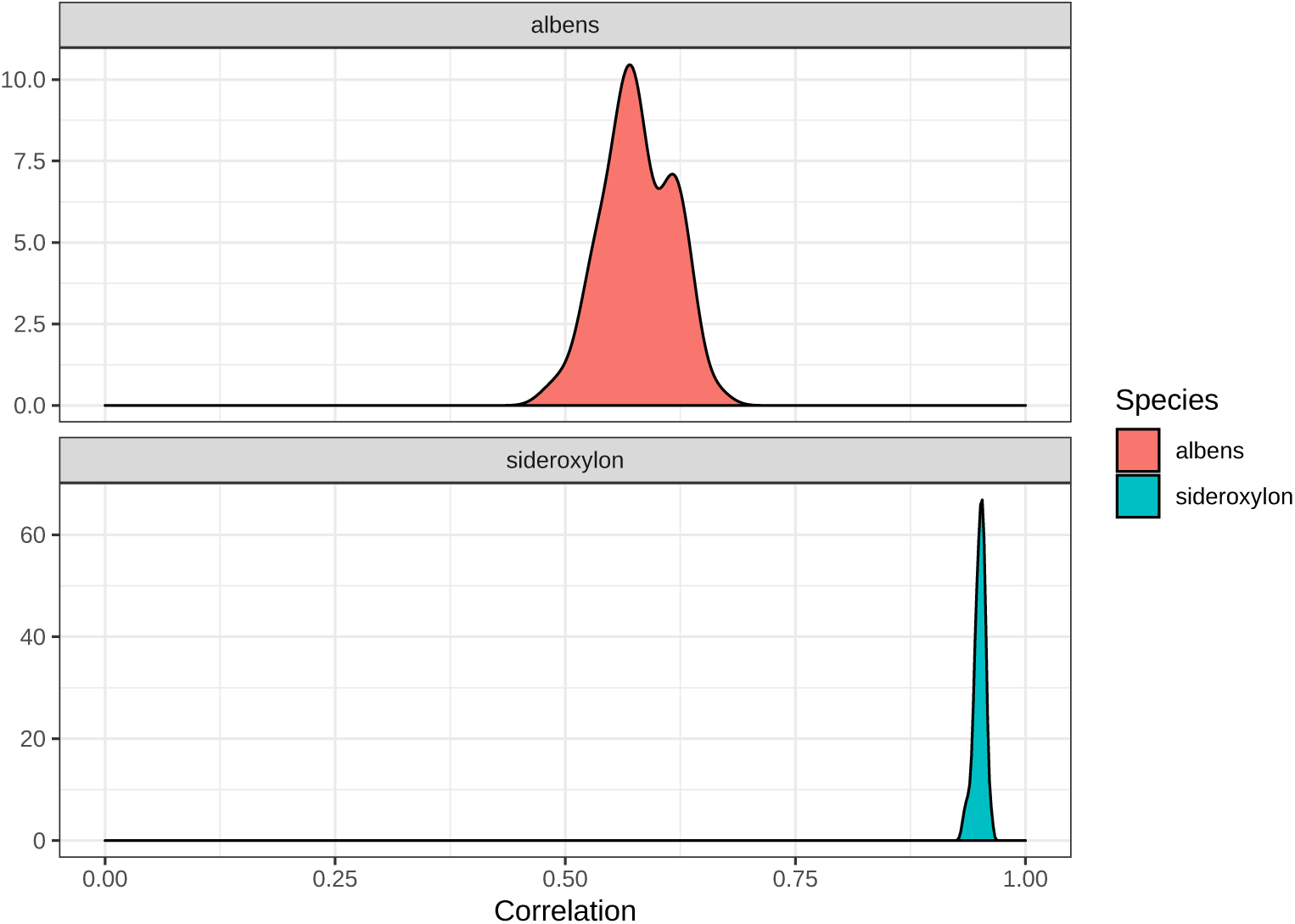
Cross-validation accuracy of best-fit GDM models for *E. albens* and *E. sideroxylon*. To test the predictive power of GDM models, GDM are fit on a training dataset with 10% of sampling locations removed in each dataset. The genetic distances of the remaining 10% of samples are predicted from their geographic and environmental data. Pearson’s correlation is used to assess the goodness-of-fit between predicted and actual genetic distances.

**Figure 11:**
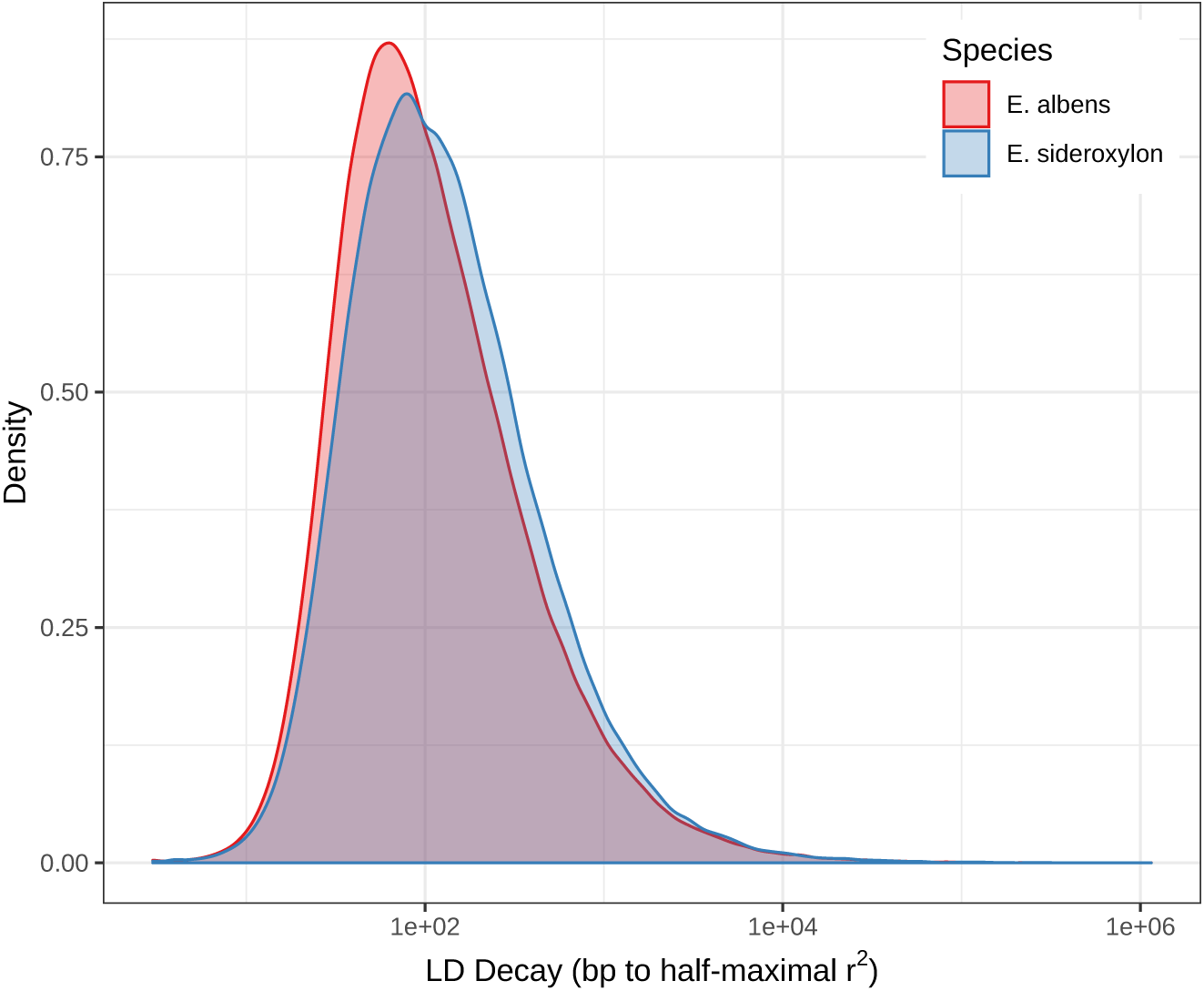
Distribution of LD extent for *E. albens* and *E. sideroxylon*. Here we show the distribution of LD extent, defined as the distance required for half-maximal decay in *R*^2^, aggregated for all 1000000 bp genome windows

**Figure 12:**
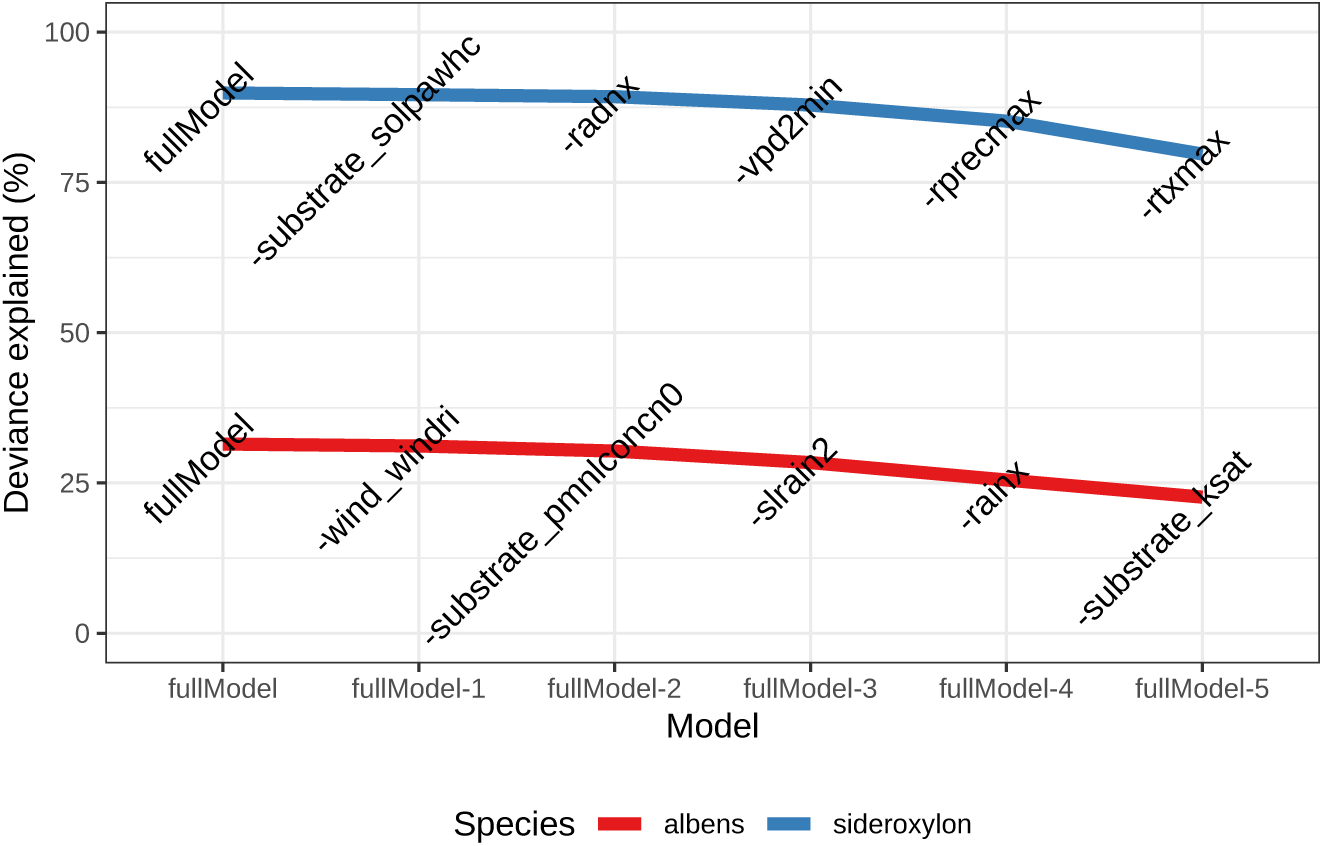
GDM model deviance explained during back-selection of variables. “fullModel” describes the model with all variables included. Each subsequent point removes one variable (per labels on plot). Please see supplementary tbl. 2 for variable names

**Figure 13:**
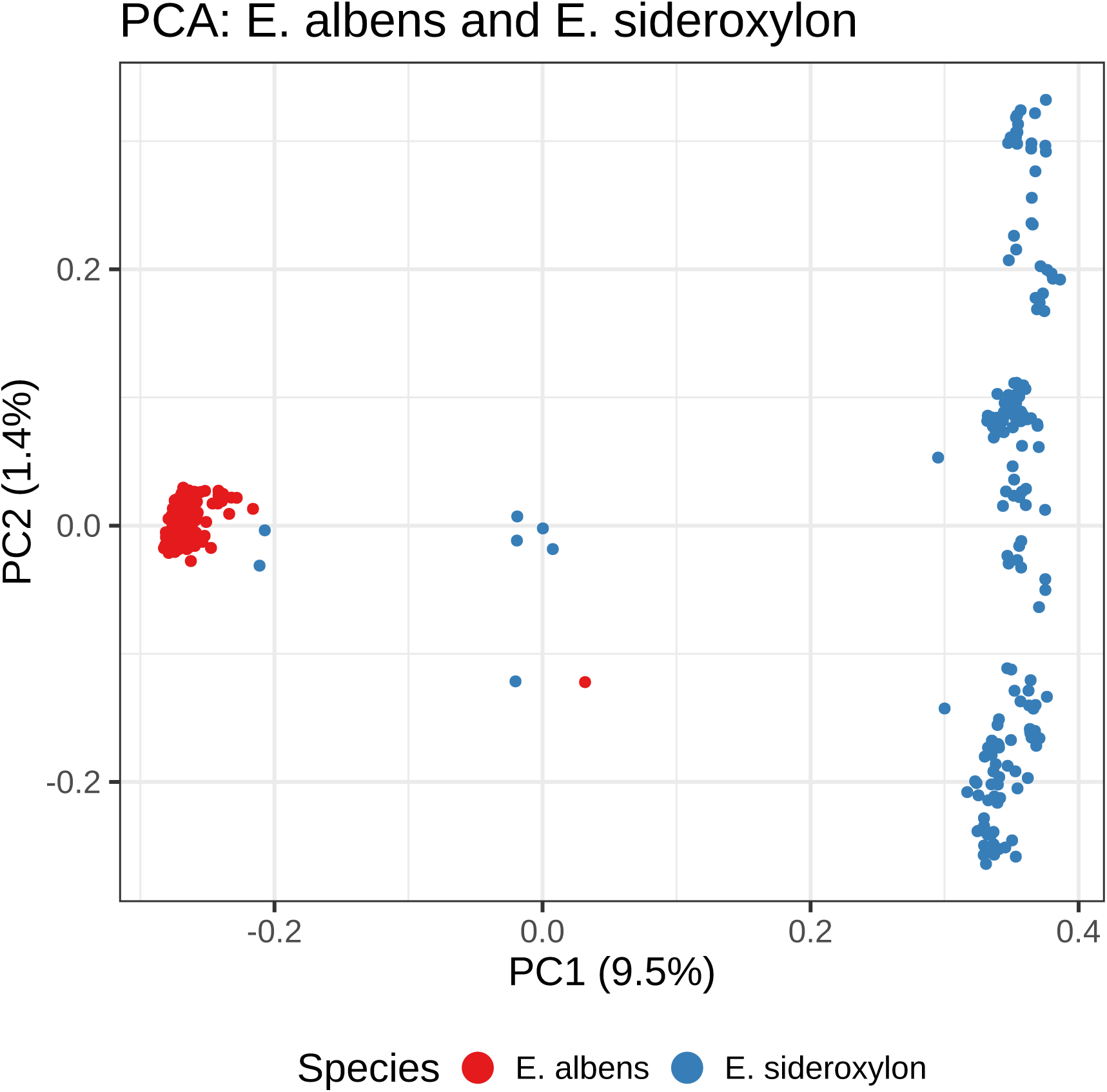
Cross-species PCA of genetic covariance estimated from PCANGSD.

**Figure 14:**
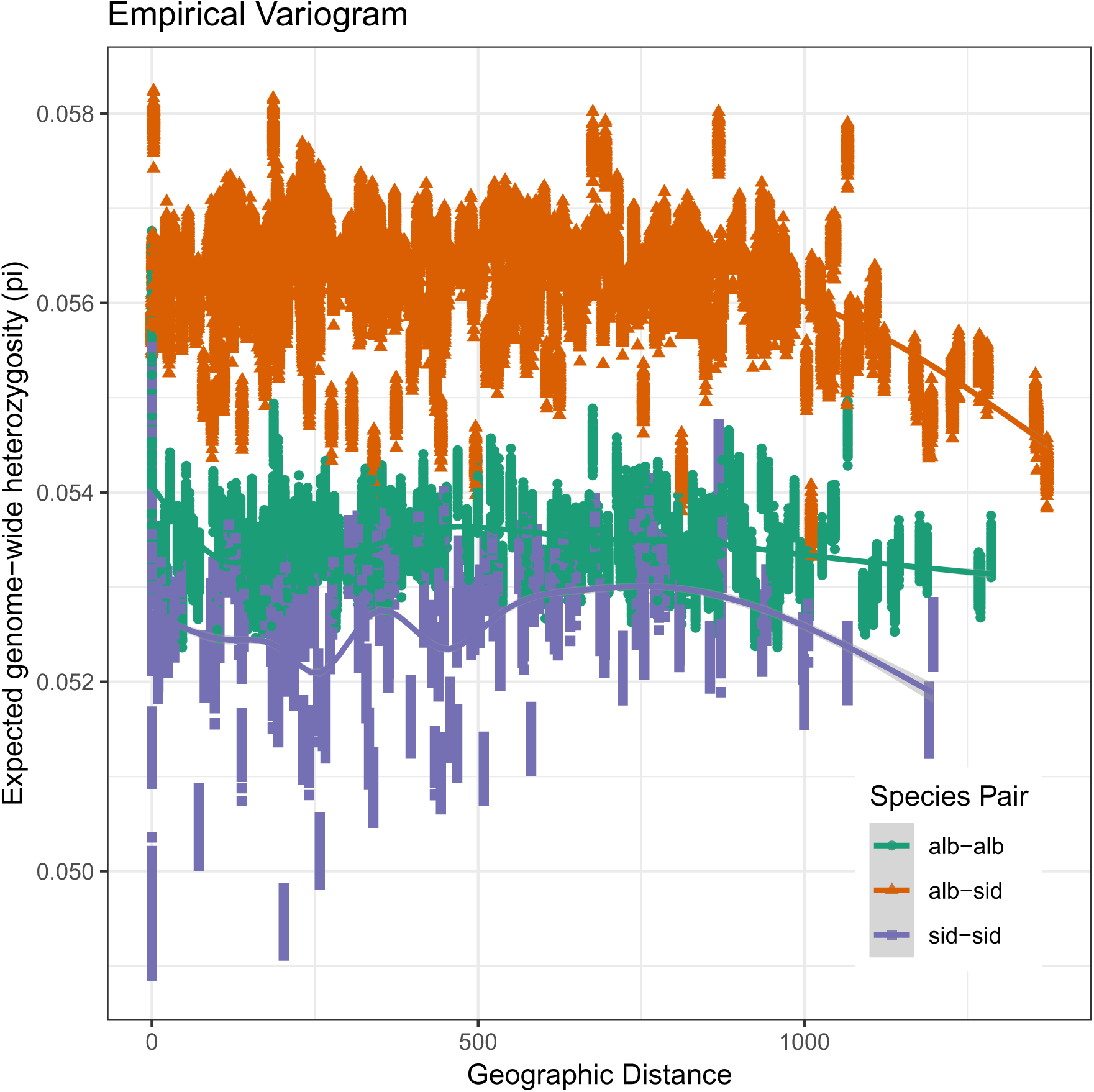
Empirical variogram of intra- and inter-species locality comparisons.

**Figure 15:**
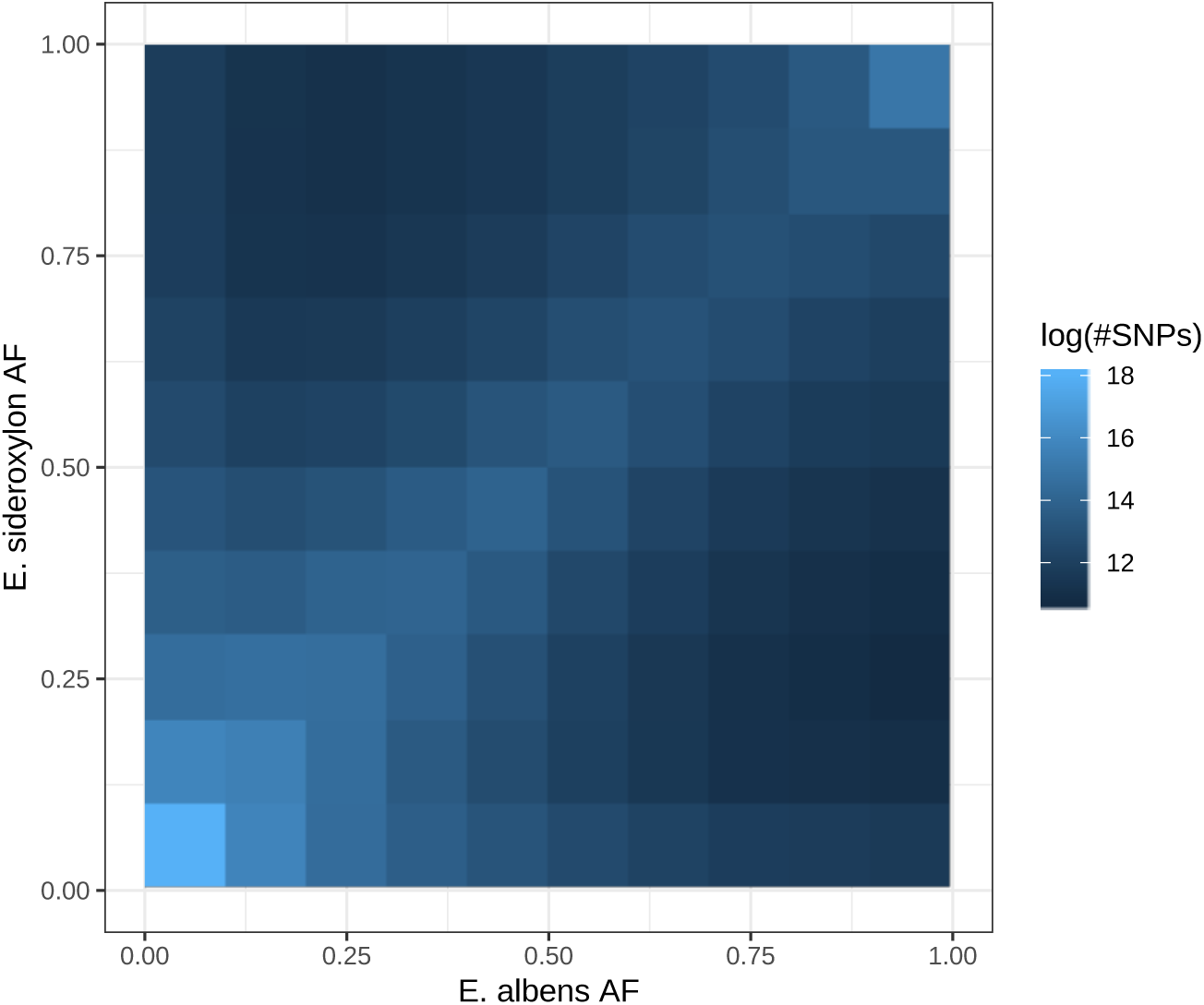
Two-dimensional site frequency spectrum between *E. albens* and *E. sideroxylon*.

**Table 3:** GDM model variables, model deviance explained, and variable-specific p-values.

|  |  |  |
| --- | --- | --- |
| *E. albens* |  |  |
| Variable | % dev. expl. | Variable p-value |
| Geographic | 31.475641 | 0 |
| Hydrologic conductivity - average saturated | 31.157494 | 0.05 |
| Precipitation - wettest month | 30.314927 | 0.02 |
| Precipitation - spring or autumn (log) season | 28.387513 | 0.13 |
| Phosphorus pre-European | 25.490866 | 0.19 |
| Wind run - month min | 22.645057 | 0.36 |
| *E. sideroxylon* |  |  |
| Variable | % dev. expl. | Variable p-value |
| Geographic | 89.887027 | 0 |
| Temperature - max difference in max | 89.599473 | 0 |
| Precipitation - max difference between successive months | 89.265338 | 0 |
| Vapour pressure deficit - month min | 87.875363 | 0 |
| Radiation - max month precipitation modified | 85.197611 | 0.04 |
| Water holding capacity - plant-available | 79.720207 | 0.12 |

**Figure 16:**
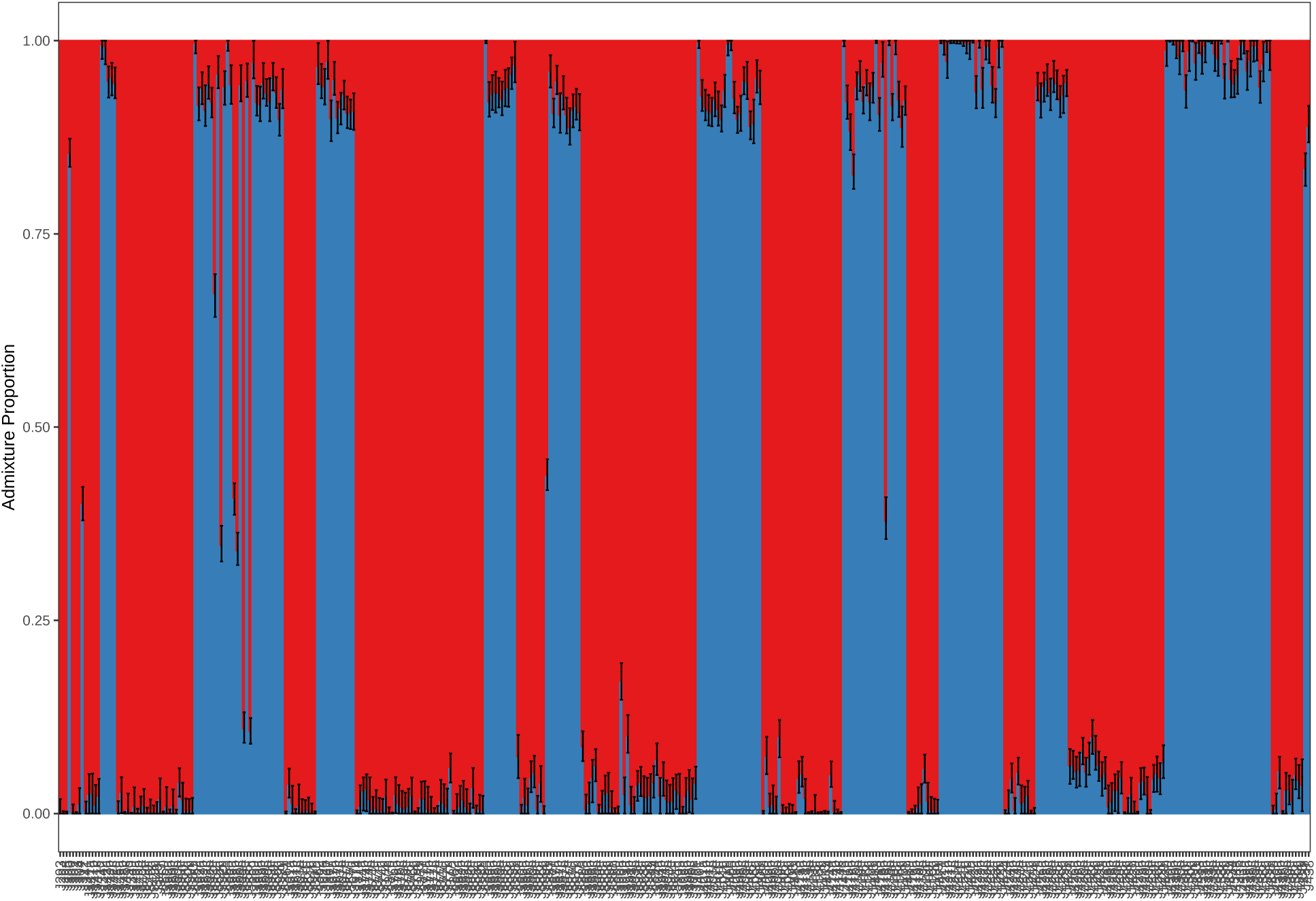
Individual-level conStruct analysis of all samples with two model layers. Admixture proportions are presented as means ± sd across 20 random subsets of 1 million SNPs. Note samples that appear as intermediates, suggesting they are recent interspecific hybrids.

